# Index-agnostic oblique plane light sheet microscopy of centimetre-scale cleared tissues at subcellular resolution

**DOI:** 10.64898/2026.06.01.729284

**Authors:** Jacob R. Lamb, Miguel Cardoso Mestre, Ksena Fenwyn Longrin, Pratiksha Bhat, Molly Stevenson, Anna D. Y. Rhodes, Joanna Gosieniecka, Leah C. Redmond, Claire A. Higgins, Noe Rodríguez-Rodríguez, Madeline A. Lancaster, James D. Manton

## Abstract

We present cleared-tissue direct-view oblique plane microscopy (CtDvOPM), which enables optically sectioned subcellular resolution imaging of centimetre-scale tissues at high throughput over the full range of clearing media refractive indices (*n =* 1.33–1.56). CtDvOPM can image conventionally-mounted expanded, aqueous or non-aqueous cleared tissue samples at up to 2 µm lateral by 14 µm axial resolution over a 10.1 mm *×* 25.0 mm *×* 6.8 mm (*x × y × z*) sample volume without image tiling, at up to 400 million voxels per second.

## Introduction

Tissue clearing, in which lipids are removed from a tissue sample and cytosol is replaced with an index-matching fluid in an attempt to homogenise refractive index and thereby reduce tissue scattering, has proven to be a powerful tool in the quest to image subcellular detail at organ-level scales [1–4]. However, the myriad tissue clearing protocols that now exist produce samples with different refractive indices as an unfortunate by-product of optimising protocols for specific tissues, antigenicity preservation and index homogeneity [5]. Furthermore, many index matching liquids are damaging to optical elements and imaging dishes, necessitating specially-designed objective lenses and sample holders.

Light sheet microscopy is well-established as a high-throughput method for imaging cleared tissues at high resolution, but currently available microscopes are limited by issues with sample mounting and restricted space-bandwidth product [6–8]. Sample mounting issues arise from the need to place two objective lenses at right angles to one another, near the sample. In many cases, this requires dipping the lenses into the (potentially corrosive) index matching fluid, or imaging through a cuvette.

Oblique plane microscopy presents an alternative approach, in which only a single objective lens needs to be placed near the specimen [9–12]. This lens is used to both excite fluorescence in a thin sheet and collect emitted light. As the light sheet is tilted with respect to the nominal focal plane, remote refocussing is used to image the tilted plane onto a camera with minimal aberrations [13]. Typically, this requires two further cofocal but non-coaxial objective lenses, and associated tube lenses.

Recently, we introduced direct-view oblique plane microscopy (DvOPM), which removes the need for the third microscope system by placing the camera directly in the remote refocussing volume [14]. Designed specifically for imaging watery samples (*n =* 1.33), our first system was capable of imaging a 370 mm^3^ volume at 2 µm by 22 µm resolution without tiling.

Here, via a re-examination of the optical theory of remote refocussing and careful imaging system design, we condense the DvOPM design down to a single lens, increase the single-tile volumetric coverage to 1700 mm^3^ and ensure the system is trivially compatible with any tissue clearing refractive index, not just that of water. We demonstrate imaging performance on samples ranging from ECi-cleared mouse gut (*n =* 1.56) to REPLICA-cleared mouse kidney (*n =* 1.33), spanning the full range of available tissue clearing refractive indices.

## Results

### Matching magnification to the sample refractive index is not optimal for remote refocussing

The goal of remote refocussing is to create a 3D image volume with minimal aberrations particularly for points away from the nominal focal plane [13, 15]. Following the method introduced by Botcherby et al., this can be achieved by matching the lateral and axial magnifications of an imaging system to the ratio of refractive indices of the remote image and sample. Such a system thereby obeys both the Abbe sine condition and the Herschel condition which, to first order, allows for stigmatic imaging of points away from the optical axis and away from the nominal focal plane, respectively. However, the theory of remote refocussing suggests that the magnification of the optical system must be changed when the refractive index is altered in order for these two conditions to be satisfied.

As much of the existing analysis of remote refocussing performance has been conducted for high numerical aperture (NA), high magnification lenses, we sought to analyse low NA, low magnification situations suitable for mesoscopic-scale imaging (detailed in Supplementary Note A). To our surprise, our investigations showed that imaging quality, as quantified by the Strehl ratio, is relatively unchanged by refractive index over ranges achievable through tissue clearing at NAs less than 0.5. This suggested that a fixed magnification set to the middle of the tissue clearing refractive index range, 1.45, would be sufficient for high quality imaging up to depths of 20 mm at an NA of 0.125, sufficient for 2 µm lateral resolution.

### Mesoscopic oblique plane microscopy using a single bi-telecentric lens

Having established that a single magnification was sufficient, we sought to maximise the field-of-view (FOV) of the optical system whilst maintaining an NA of at least 0.125. Our previous DvOPM system already featured the largest such FOV possible (6.25 mm) with traditional microscope objective lenses (using a 4*×* 0.16 NA lens) and so we instead turned to consider lenses used for optical metrology. In particular, a wide range of bi-telecentric lenses designed for parts inspection are commercially available with large FOVs and suitable NAs. Furthermore, as these lenses are bi-telecentric, a single monolithic lens can be used for the entire optical system, effectively compressing two objective lenses and two pupil relay lenses into a compact one-lens system that is inherently aligned as part of the manufacturing process. We selected a 0.69*×* bi-telecentric lens supporting an f-number of f/4 which, when used in reverse, corresponds to a lateral magnification of 1.45*×* at an NA of 0.125 over a 17.6 mm diameter FOV.

Due to refraction at the coverslip-sample interface, a light sheet launched at a fixed angle in air will have a sample refractive index-dependent angle in the sample volume (for details, see Supplementary Note B). Using a separate light sheet launch gimballed around the focal point of the selected detection lens (Figure 1d), such that the angle of incidence can be trivially varied to ensure the light sheet remains coplanar with the detection plane of the camera irrespective of refractive index (Figure 1e), we could illuminate with a Gaussian light sheet the full FOV of our selected camera, 10.1 mm by 6.8 mm, corresponding to a 300 % increase over our previous DvOPM system.

**Figure 1.**
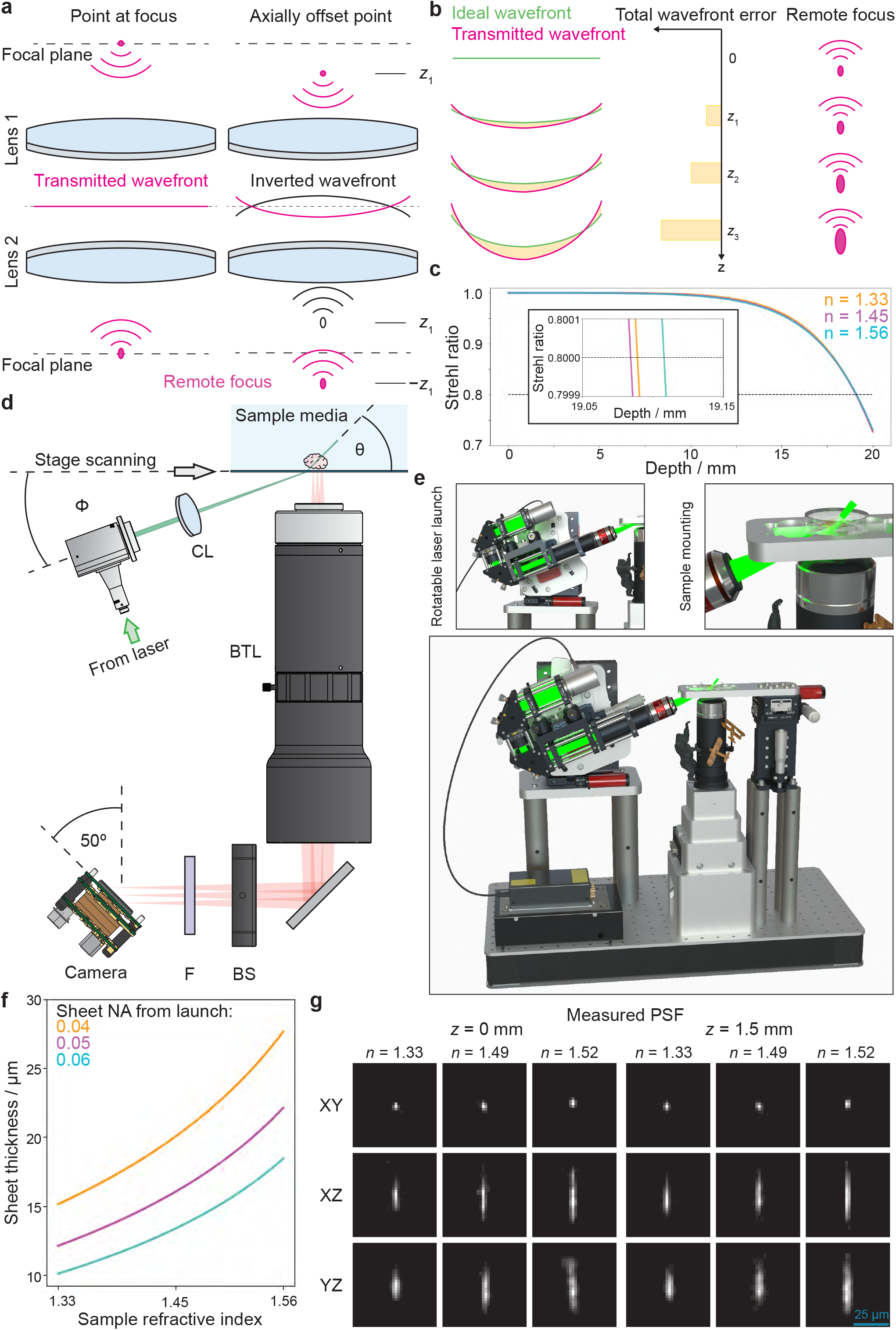
Cleared-tissue direct-view oblique plane microscopy concept and implementation. **a** Illustration of remote refocus imaging of displaced point showing transmitted wave in red and the ideal refocussing wavefront in black. As a result of the mismatch between these two wavefronts, a point closer to lens 1 is imaged to a point further away from lens 2, rather than closer. **b** Illustrative comparison of ideal and transmitted wavefronts for the remote refocussing scenario shown in a. As imaging depth increases, the total wavefront error, shown in yellow, grows and leads to an increasingly aberrated remote focus (red). **c** Calculated Strehl ratios as a function of depth for a fixed magnification and varying refractive index. For all refractive indices, diffraction-limited performance (Strehl ratio > 0.8) is maintained over a depth of more than 19 mm (inset). **d** Optical diagram of CtDvOPM implementation, including bi-telecentric lens (BTL), beam shifter (BS), filter (F), cylindrical lens (CL) and angle-adjustable (*φ*) light sheet launch. Volumes are acquired by scanning the sample through a static light sheet using a motorised stage. **e** CAD rendering of implemented CtDvOPM instrument. Insets highlight the angle-adjustable light sheet launch with refractive-index selection chocks (red), and sample mounting (compatible with 35 mm imaging dishes and multiwell slides). **f** Plot of sheet thickness along *z* axis as a function of refractive index for varying sheet launch numerical apertures (NAs). For all NAs, the thinnest sheet is achieved at the lowest refractive index. **g** Measured point spread functions (PSFs) obtained using sub-diffraction beads showing diffraction-limited performance for all refractive indices, *n*, and depths, *z*, of 0 mm and 1.5 mm.

### Light sheet thickness decreases with decreasing refractive index

As the light sheet is created in air, passes through a glass coverslip and then refracts at the interface between the glass and the sample, changing refractive index alters the angle of refraction. While this is compensated for by changing the angle of the light sheet launch, this compensation is only exact for the chief ray of the light sheet. As the sheet is formed from waves spanning a small range of angles, each refractive index leads to a slightly different compression of this angular range. Figure 1f shows the result of this, displaying the thickness of the light sheet as a function of sample refractive index. Perhaps surprisingly, the thinnest light sheet is achieved in the lowest refractive index (*n =* 1.33). This is discussed further in Supplementary Note C.

### CtDvOPM achieves Nyquist-limited resolution for all relevant refractive indices

By using sub-diffraction beads, we acquired point spread functions (PSFs) in water (*n =* 1.33), OptiMuS RI-matching solution (*n =* 1.49) and immersion oil (*n =* 1.52), shown in Figure 1g. Gaussian fitting of at least 10 beads per condition produced full-width half-maxima of (2.7 *±* 0.4) µm, (3.1 *±* 0.3) µm and (2.8 *±* 0.2) µm across the untilted axis and (2.8 *±* 0.4) µm, (3.3 *±* 0.3) µm and (2.6 *±* 0.3) µm across the tilted axis. As expected from the theoretical plots in Figure 1f, the watery PSF had the smallest axial extent of (13.6*±*2.9) µm, compared with (18.5*±*1.8) µm for OptiMuS RI and (22.9*±*2.3) µm for immersion oil.

### CtDvOPM provides organ-scale imaging at sub-cellular resolution for all relevant refractive indices

Encouraged by our PSF measurements, we first imaged a mouse gut cleared using ECi, with a refractive index of 1.56 (Figure 2a) [3]. Here we were able to clearly resolve the internal structure of individual villi. Next, we rotated the light sheet launch to the calculated angle for *n =* 1.33 and imaged a watery REPLICA-cleared kidney slice sample (Figure 2b). Apart from a small refocussing of the light sheet, no further alignment was required to produce clear images of the sample despite the drastic change in refractive index (∆*n =* 0.23). Here, image quality was sufficient to clearly see individual collecting ducts, demarked by aquaporin 2, over the entire 500 µm thickness of the tissue slice.

**Figure 2.**
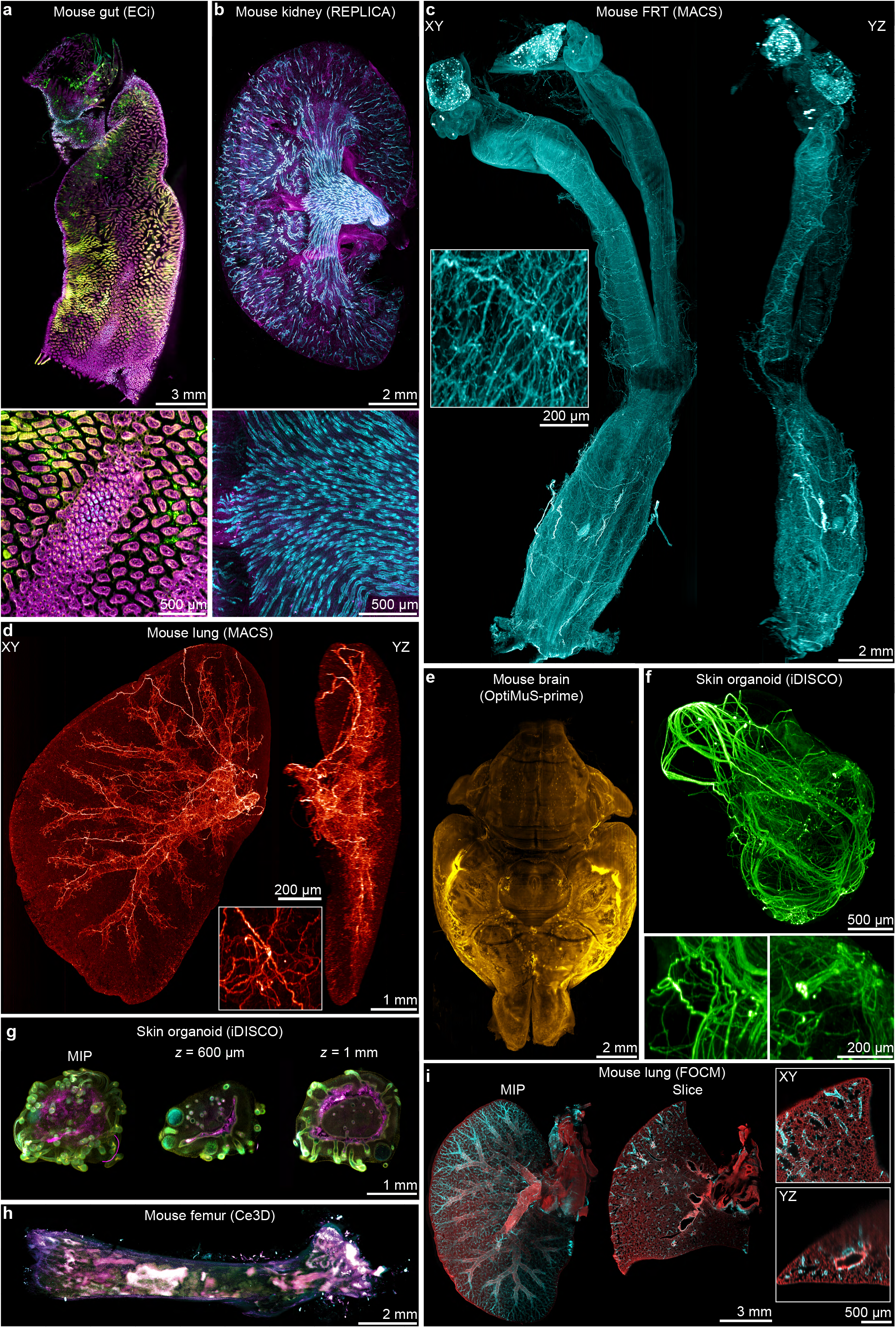
Direct-view oblique plane microscopy can image cleared tissues with any existing refractive index (n = 1.33–1.56). **a** Multicolour single laboratory-frame XY slice of a mouse gut cleared using the ECi protocol (*n =* 1.56). Labelled nuclei are shown in magenta and tuft cells in green. **b** Maximum intensity projection (MIP) of a 500 µm-thick mouse kidney slice cleared using the REPLICA protocol (*n =* 1.33). Immunolabelled aquaporin 2 signal, highlighting collecting ducts, is shown in cyan and immunolabelled CD105 is shown in magenta. **c** MIP of mouse female reproductive tract (FRT) immunolabelled for PGP9.5 and cleared using the MACS tissue clearing kit (*n =* 1.56). Inset shows zoom of a highly innervated region. **d** MIPs of mouse lung lobe immunolabelled for beta III tubulin and cleared using the MACS tissue clearing kit (*n =* 1.56). **e** Adult mouse brain cleared using OptiMuS-prime (*n =* 1.49) and immunolabelled for beta III tubulin. Staining in this sample was largely nonspecific. **f** Induced pluripotent stem cell (iPSC)-derived skin organoid (chimpanzee) cleared using the iDISCO protocol (*n =* 1.56) and immunolabelled for NFH (neurofilament heavy), showing myelinated peripheral neurons. **g** iPSC-derived skin organoid (human) cleared using the iDISCO protocol (*n =* 1.56) and immunolabelled for Ki-67 (green) showing proliferating cells, caspase-3 (magenta) indicating apoptotic cells and Hoechst 33258 (cyan) labelling nuclei. **h** Mouse femur expressing BFP (blue) and tdTomato (magenta) immunolabelled with a neuronal marker (yellow) and cleared using the Ce3D tissue clearing kit (*n =* 1.50). **i** Mouse lung lobe immunolabelled for *α*-actin (cyan) and with strong autofluorescence signal (red), cleared using the FOCM protocol (*n =* 1.495). Note: all images have been heavily downsampled and compressed to comply with bioRxiv’s 40 MB filesize limit. Full-size images are available to view at www.ctdvopm.org.

We then imaged an entire mouse female reproductive tract cleared using the commercial MACS workflow (*n =* 1.56) as a further test of the compatibility across tissue clearing approaches. Here, neuronal processes could be seen traversing the entire 25 mm length of the tissue. We also separately segmented bronchi and neurons over the over the 130 mm^3^ volume of a MACS-cleared mouse lung (Figure 2d, Figure 3).

**Figure 3.**
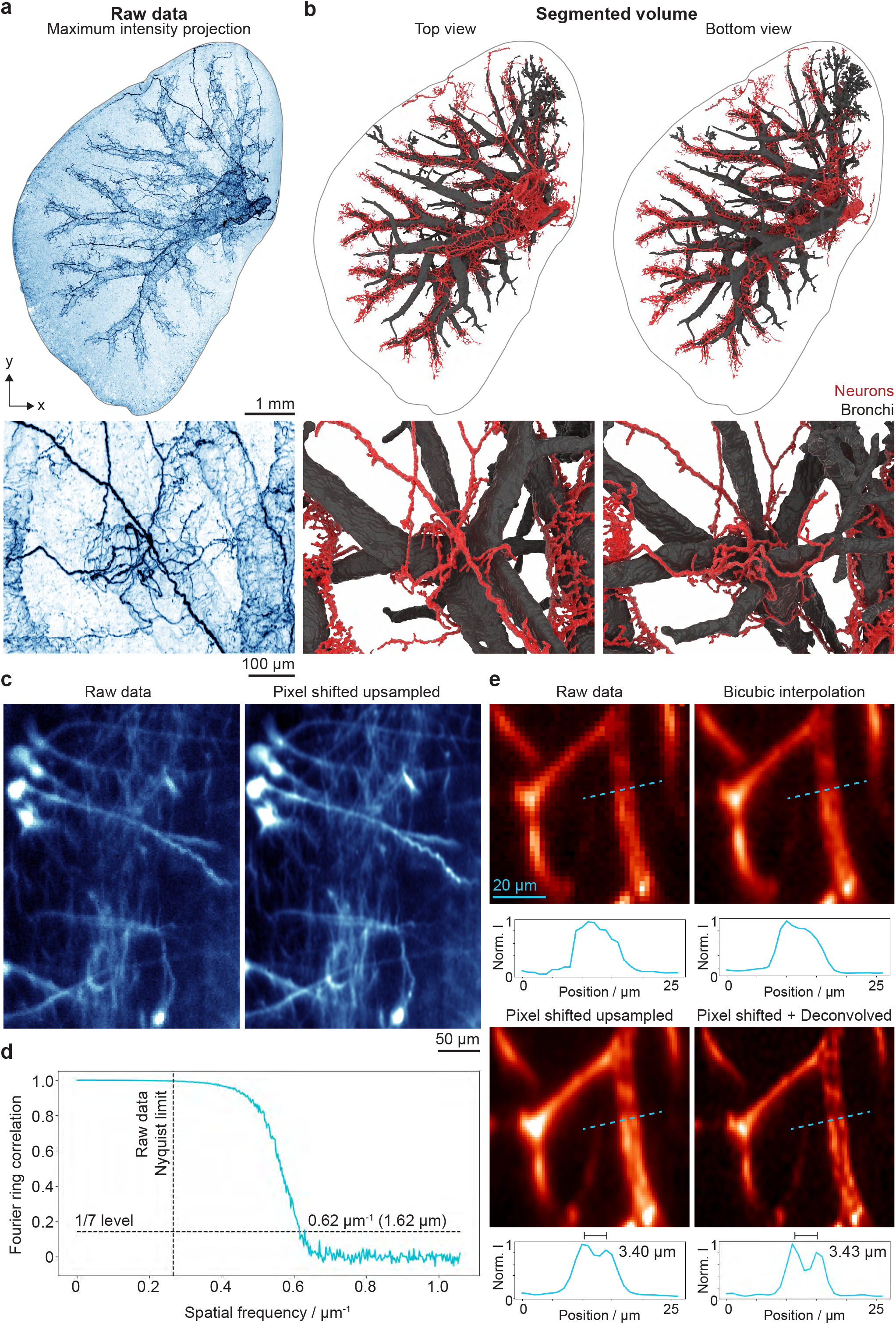
Image processing using cleared-tissue direct-view oblique plane microscopy data. **a** Maximum intensity projection of mouse lung lobe data shown in Figure 2d. **b** 3D surface render of automatically segmented, flood-filled volumes for bronchi (grey) and neurons (red). Below each panel zoomed views of the corresponding datasets in a and b are shown. Fine neuronal processes can be seen wrapping around a branch point in the bronchial network from both above and below. **c** Comparison between the raw (undersampled) and reconstructed pixel shifted upsampled data of a subregion of a SmartBatch+ clear mouse brain, illustrating the improvement in image fidelity. Extended views of this sample can be found in Supplementary Figure 6. **d** Fourier ring correlation plot of the pixel shifted upsampled data, shown in c, clearly demonstrating spatial frequency correlations extending beyond the Nyquist sampling limit of the raw data. **e** Comparison between raw, bicubic interpolated, pixel shifted upsampled and deconvolved pixel shifted upsampled data for a subregion of a CUBIC-cleared mouse cerebellum. Line profiles (shown in blue) through an apparent single filament show that only in data that has been pixel shifted can this structure be accurately resolved as two separate filaments. These are separated by 3.4 µm, less than the 3.8 µm Nyquist sampling limit of the raw data. A fuller view of this sample can be found in Supplementary Figure 7.

The 10 mm-wide FOV of the system was sufficient to image an entire adult mouse brain without tiling (Figure 2e, cleared using OptiMuS-prime, *n =* 1.49) and could, using a recently developed aliasing approach to increase acquisition speed, theoretically be completed in less than a minute given a sufficiently bright sample [16]. Smaller specimens, such as the ∼2 mm diameter hair organoid shown in Figure 2f, can be acquired even faster, with simple camera region-of-interest selection increasing the potential maximum frame rate. With a motorised filter wheel, this acquisition mode could support full volumetric multicolour imaging in less than 1 min (Figure 2g).

As a further test of tissue clearing protocol compatibility, we imaged a Ce3D-cleared mouse femur (Figure 2h) and mouse lung cleared using the rapid FOCM protocol (Figure 2i) [4, 17]. While the quality of tissue clearing in both of these samples was markedly less than in the others, gross anatomical features could still clearly be seen.

### Sub-pixel image shifting extends resolution beyond the Nyquist sampling limit

Given the physical size of pixels on the camera chip, the lateral resolution is limited by Nyquist sampling to 3.8 µm. To circumvent this, we integrated an image shifter into the optical system to provide sub-pixel image shifts through which a higher resolution (2 µm, at the diffraction limit of the 0.125 NA lens used in this work) can be reconstructed. Figure 3c shows both a raw image and upsampled image calculated from a 2*×*2 image shift grid, with the upsampled image showing markedly improved resolution and contrast. This is quantified by the Fourier ring correlation plot shown in Figure 3d, where the 1/7 threshold is reached at twice the raw data Nyquist limit. Furthermore, Figure 3e shows that merely in-terpolating the raw data into an upsampled grid does not lead to an increased resolution, as demonstrated by the inability to distinguish a gap between two parallel filaments. In contrast, the image-shifted upsampled data shows a clear delineation that is further enhanced by deconvolution.

## Discussion

Here we have introduced cleared-tissue direct-view oblique plane microscopy as an index-agnostic instrument for rapid, volumetric cleared-tissue imaging at subcellular resolution using conventionally mounted samples. By default, the lateral resolution is limited by Nyquist sampling to 3.8 µm although, by integration of an image shifter into the optical system to provide sub-pixel image shifts, a higher resolution image (2.0 µm resolution, at the diffraction limit of the 0.125 NA lens used in this work) can be reconstructed. Axial resolution is limited by the need to keep the light sheet sufficiently thin over the full FOV and so a resolution better than the current 13.6 µm is not possible with a static sheet. Unfortunately, the nature of refraction at the sample interface precludes the easy integration of axially swept light sheet microscopy, in which a much thinner but shorter sheet is swept axially in tandem with the rolling shutter of the camera (see Supplementary Note C for details) [18]. Nevertheless, the current resolution is sufficient for subcellular visualisation and neuronal tracing, with the rapid imaging speed and conventional sample mounting providing an avenue towards automated high-throughput volumetric imaging of hundreds of samples.

While the bi-telecentric lens used here already surpasses the FOV of conventional objective lenses with comparable resolution, there is scope to extend the FOV by a further order of magnitude by the selection of alternative bi-telecentric lenses and cameras. While lateral FOV can always be computationally extended by image tiling, the axial extent of the imageable volume is limited by the working distance of the lens used. Here, the use of bi-telecentric lenses provides a further advantage over conventional microscope objective lenses as their working distances are significantly greater.

Together, our theoretical analysis and experimental data show that direct-view oblique plane microscopy enables straightforward cleared-tissue imaging across refractive indices using conventional sample mounting, with resolu-tion and FOV matching or exceeding existing light-sheet approaches and without the light loss associated with non-direct mesoscopic oblique plane microscopes. The implementation described herein costs less than 45 k€ and can be assembled in less than a day. We have provided open source hardware and software at https://www.ctdvopm.org and are committed to helping other researchers implement such systems within their laboratories. We anticipate that this implementation of direct-view oblique plane microscopy will provide both a practical platform for volumetric cleared-tissue imaging and an extensible basis for further improvements in imaging performance.

## Acknowledgements

JRL and JDM thank Adam Fowle and Elliot Oakman of the LMB’s Mechanical Workshop for fabricating the custom optomechanics used in this work, and Rebecca McClelland for proofreading the manuscript. We are grateful to the Biological Service Group and Ares staff, and the genotyping facility for their technical assistance and support. JDM acknowledges support from the Royal Society through a University Research Fellowship (URF/R1/221086 and RF/ERE/221078). MS, NR-R and MAL acknowledge support from the Medical Research Council as part of United Kingdom Research and Innovation (U105178805, MC_UP_1201/35, MC_UP_1201/9). NR-R has grant funding from the European Union’s Horizon 2020 research and innovation program under the Marie Skłodowska-Curie grant agreement number 896454. PB acknowledges support through a research collaboration between AstraZeneca UK Limited and the Medical Research Council (Blue Sky programme). CAH acknowledges support from the Biotechnology and Biological Sciences Research Council in collaboration with Procter & Gamble (BB/X511249/1), and the LEO Foundation.

## Data availability

As the datasets generated by CtDvOPM are large, we have made a subset of the raw and processed datasets generated in this work available online, while others will be provided direct from the authors upon request. MACS-cleared lung and REPLICA-cleared kidney datasets are available from https://doi.org/10.5281/zenodo.20489678 and https://doi.org/10.5281/zenodo.20490386, respectively. An exemplar dataset of the image shifting approach shown in Figure 3e is available at https://zenodo.org/records/20828379. Full CAD files used to construct a CtDvOPM instrument can be downloaded from https://zenodo.org/records/20828373.

## Methods

### Cleared-tissue direct-view oblique plane microscope design and implementation

The microscope detection system consists of a single 0.69*×* CobaltTL f/4 telecentric lens (Edmund Optics # 15–872). The lens is mounted in reverse, producing a bi-telecentric imaging system with a magnification of 1.45 and an object-side numerical aperture of 0.125. Fluorescence light collected by the lens is focused through an emission filter to remove reflected laser light before being imaged directly onto an IDS U3-31R2SE Rev.1.2 camera (IMX540 sensor). To minimise aberrations, the camera cover glass was removed by EURECA Messtechnik GmbH. The lens, filter wheel, and camera are mounted using a combination of adapted Thorlabs components and custom parts. These components were designed such that the entire detection system can be assembled without the need for optical alignment. The spacing between components was optimised using OpticStudio Zemax simulations with black-box lens files provided by Edmund Optics.

Excitation light is generated by a fibre-coupled four-channel laser (Cobolt Skyra 405/488/561/638) coupled into a 30 mm cage system using a reflective fibre collimator (Thorlabs RC12APC-P01). The beam then passes through a 100 mm focal length cylindrical lens (Thorlabs ACY256-100-A) and a long working distance 2*×* super apochromatic microscope objective (Thorlabs TL2X-SAP) to generate the light sheet within the sample. The objective lens is used to generate the sheet in order to minimise wavelength-dependent focal shifts during multicolour imaging. As shown in Figure 1e, the cage system is mounted on a custom rotatable plate, allowing the light sheet to remain coplanar with the camera across dif-ferent sample refractive indices. The laser launch is designed to rotate about the point where the light sheet intersects the cover glass. Alignment to the correct angle for a given refractive index is achieved using custom 3D-printed chocks inserted into the rotating assembly. Fine positioning of the light sheet is achieved by laterally translating the entire laser launch parallel to the optical bench, while adjustment of the sheet waist position is performed using a manual translation stage aligned with the optical axis of the excitation lens.

The microscope uses a custom sample stage compatible with both 35 mm dishes and multi-well plates. Volumetric acquisition is achieved by translating the sample through a static light sheet using a motorised translation stage (Thorlabs ZFS25B).

The microscope is controlled using Micro-Manager, while a custom Python GUI is used to control the position of the light-sheet focus stage. A complete set of CAD files for the CtDvOPM is available via https://www.ctdvopm.org, which also features an interactive CAD render and a full parts list (also provided in Supplementary Note D).

### Image processing

#### Image volume deskewing and rotation

Raw data are deskewed into volumes rendered in the laboratory coordinate frame using an affine transformation implemented in a custom Python script. Full details on this can be found in Supplementary Note E. All non-image-shifted data presented is acquired using a slice spacing of 5 *µ*m and when deskewed has an isotropic pixel size of 1.884 *µ*m in the laboratory coordinate frame.

#### Image-shifter data upsampling

For diffraction-limited imaging beyond the Nyquist limit imposed by the 2.73 µm physical size of the pixels on the camera employed in this work, we integrated an Optotune BSW-20-4.8-VIS-M beam shifter into the optical system before the filter wheel. After calibration using fluorescent beads, we applied voltages to the ICC-4C-2000 controller such that the image was shifted by (0 µm, 0 µm), (0 µm, 1.046 µm), (1.365 µm, 0 µm) and (1.365 µm, 1.046 µm), recording an image for each of these half-pixel shift combinations (the shifts do not have the same magnitude in the two axes due to the 50^*°*^ tilt of the camera sensor normal with respect to the optical axis.).

We found that the standard ‘shift-and-add’ algorithm used to calculate upscaled data from sub-pixel shifts led to a low visibility of fine features, as low spatial frequencies are over-represented in the reconstruction (i.e. the effective modulation transfer function is damped at higher spatial frequencies). As such, we repurposed the Richardson–Lucy algorithm to develop an iterative method of reconstructing diffraction-limited data without this issue. Briefly, the ground truth object estimate was formed on a 2 *×* 2 upsampled grid and blurred with a theoretical PSF calculated on the same grid. Four copies of this image were created, with each one shifted by an amount corresponding to one of the raw captured images and then binned 2*×*2, forming the four predicted shifted images. These were then compared with the raw captured images using elementwise division to form an error map, with an update map being formed by splatting each of the error maps 2 *×* 2, shifting in the opposite direction to that in the forwards operator, summing the four arrays elementwise and blurring with a flipped PSF. We implemented the gradient consensus approach to implicit regularisation to ensure that the iterations terminated at an appropriate point, with the final upsampled image being created from convolving the final estimated ground truth object with the theoretical point spread function. In this way, any deviations between the real point spread function and the theoretical one used are made irrelevant and any increase in resolution must have come from the upsampling and not the deconvolution step. Fourier ring correlations were calculated using the one-image approach of Rieger et al. [19].

### Tissue sample preparation

Antibody details for all samples are presented in Table 1.

**Table 1.**
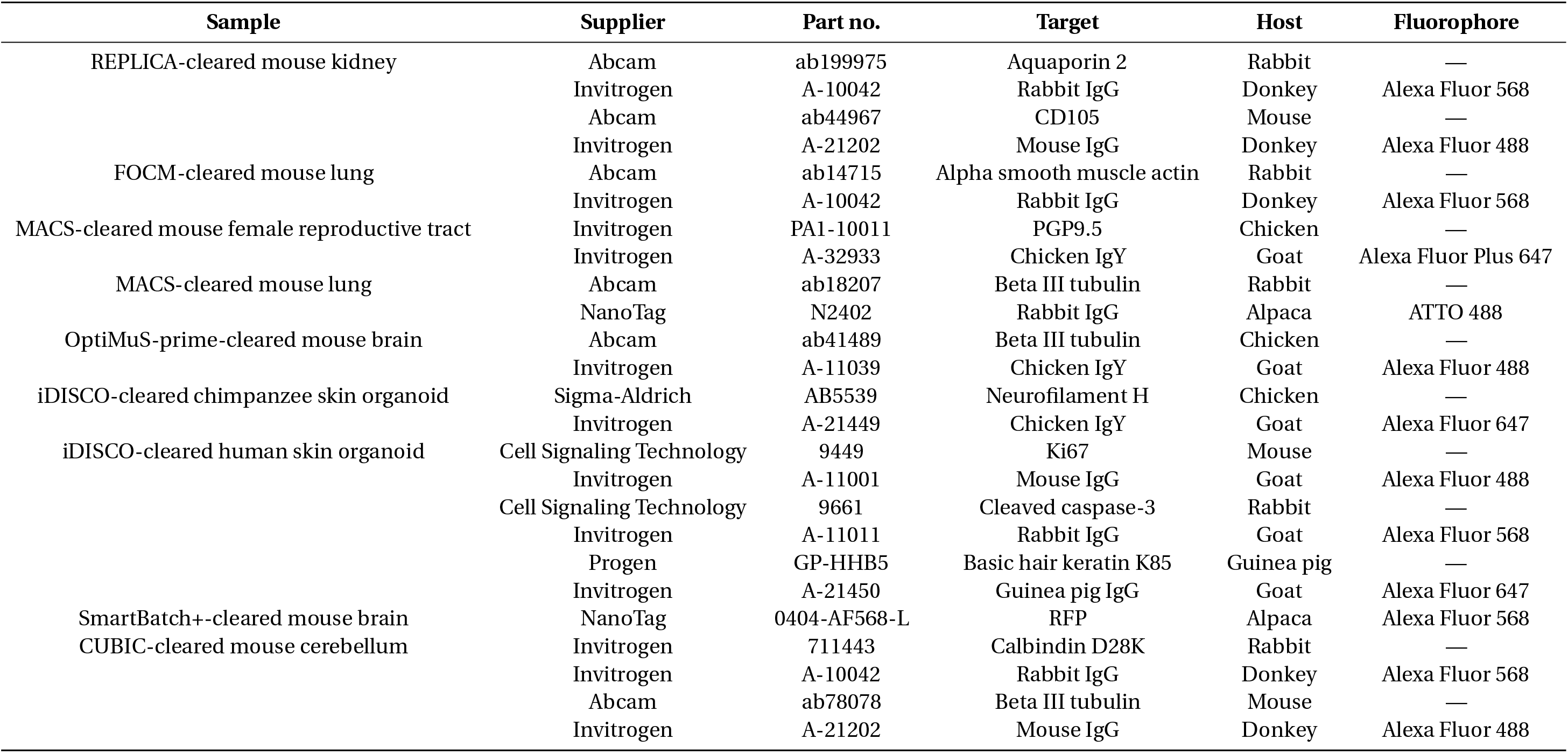
Antibody details for cleared tissue samples.

#### iDISCO tissue clearing (skin organoids)

The iDISCO protocol used to stain the 3D organoids was produced by combining the original published iDISCO protocol with the one written specifically for hairy skin organoids [2, 20]. The protocol was 17 days long, as explained here:

Day 1: Organoids were collected and washed twice with 1X PBS (Merck, D8537) (2 ml) to remove any residual medium. Organoids were fixed in 4% paraformaldehyde (PFA) (Lifetech, 2890) in a full 2 ml Eppendorf tube for 24 hours at 4 °C on a tube rotator.

Day 2: Organoids were washed at room temperature with shaking 3 times in PBS (2 ml) for 30 minutes, 1 hour, and 1 hour respectively to remove PFA. After washing, organoids were stored in PBS with 0.02 % sodium azide (Sigma, S2002) at 4 °C to prevent bacterial growth.

Day 3: Fixed organoids were washed in 2 ml PTx.2 at room temperature for an hour, twice. PTx.2 (1 litre) is composed of 100 ml PBS 10X (Thermo, 14190250), 2 ml Triton X-100 (Thermo, HFH10), 898 ml Milli-Q water. Organoids were then incubated in 2 ml 1 *×*PBS/ 0.2 % Triton X-100/ 20 % DMSO (Sigma, D8418-50ML), 37 °C, overnight.

Day 4: Organoids were incubated in 2 ml 1*×*PBS/ 0.1 % Tween-20 (Sigma, P2287)/ 0.1 % Triton X-100/ 0.1 % sodium deoxycholate (Sigma, D6750-100G)/ 0.1 % NP40 (Sigma, 492016-100ML)/ 20 % DMSO at 37 °C overnight.

Day 5: Organoids were washed in 2 ml PTx.2 twice for one hour each at room temperature. They were then incubated in 2 ml permeabilisation solution at 37 °C for 24 hours. Permeabilisation solution (50 ml) is composed of 40 ml PTx.2, 1.5 g of glycine (Sigma, G7126-1KG) and 10 ml of DMSO. Permeablisation solution was made on the day, due to glycine’s tendency to precipitate out of solution at low temperatures.

Day 6: Organoids were blocked in 2 ml blocking solution for 24 hours at 37 °C. Blocking solution (50 ml) is composed of 42 ml PTx.2, 3 ml of goat serum (Abcam, ab7481-50ml), 5 ml of DMSO.

Days 7–9: Organoids were first transferred from 2 ml Eppendorf to 200 µl PCR tubes in order to reduce the volume of primary antibody required. Primary antibodies were diluted to their appropriate concentrations in PTwH / 5 % DMSO / 3 % goat serum. Organoids were then incubated in 200 µl of their primary antibodies for 67 hours at 37 °C. PTwH (1 litre) was made up of 100 ml PBS 10X, 2 ml Tween-20, 1 ml of 10 mg/ml heparin stock solution (Sigma, H3149-50KU), 898 ml Milli-Q water.

Day 10: Organoids were transferred back to their original 2 ml Eppendorfs, and washed in 2 ml PTwH 4–5 times at room temperature over the space of 24 hours.

Day 11–14: Organoids were transferred to a fresh set of 200 µl PCR tubes and incubated with secondary antibody diluted in PTwH / 3 % goat serum, 37 °C, 67 hours. Secondary antibodies were filtered using a 20 µm sterile filter before being applied to the organoids to prevent formation of precipitates in the sample.

Day 15: Organoids were transferred back to their original 2 ml Eppendorfs, and washed in 2 ml PTwH for 4–5 times at room temperature over 24 hours.

Day 16: Organoids were incubated with 2 ml Hoechst (Thermo, H3569) diluted in PBS (at final concentration of 120 µg/ml) for 3 hours at room temperature. Organoids were then dehydrated in a methanol (VWR, 83638.320) / H_2_O series: 20 %, 40 %, 60 %, 80 %, 100 %, 100 %; 2 ml, 1 hour each at room temperature.

Day 17: 100 % methanol was removed from the organoids, and replaced with 2 ml 66 % DCM (Sigma 270997-12X100ML) / 33 % methanol in the Eppendorf. This was left at room temperature, with shaking, for 3 hours. This liquid was then replaced with 100 % DCM for 15 minutes at room temperature with shaking, twice over, to remove all methanol from the sample. Organoids were then incubated in 2 ml dibenzylether (DBE, Sigma 108014-1KG) (no shaking) for 2 hours at room temperature, and stored indefinitely at 4 °C.

#### MACS tissue clearing (mouse lung and FRT)

Mouse lung and female reproductive tract (FRT) tissues were processed for whole-mount immunostaining and optical clearing using the MACS Clearing Kit (Miltenyi, 130126-719) protocol. Freshly dissected lungs were collected in PBS buffer, fixed overnight at 4 °C in freshly prepared 4 % formaldehyde in PBS (ThermoFisher, 28908), and washed for 24 hours with three PBS buffer changes; if not processed immediately, samples were stored in PBS/azide buffer at 4 °C. Each lung was permeabilised in 5 ml MACS Permeabilization Solution for at least 24 hours at room temperature under slow continuous rotation, after which samples were transferred to 1.5 ml freshly prepared 1*×* MACS Antibody Staining Solution containing either a primary antibody or a primary conjugated to a smart secondary (NanoTag) in a 24-well plate. Staining was performed protected from light under slow continuous rotation for 7 days at 4 °C. Samples were then washed three times in 5 ml 1*×* MACS Antibody Staining Solution under slow continuous rotation at room temperature, consisting of two four-hour washes followed by an overnight wash. When secondary antibodies were used, samples were incubated in 1.5 ml 1*×* Antibody Staining Solution containing the secondary antibody under the same light-protected shaking conditions for 7 days at 4 °C, followed by the same three-step washing procedure. Tissues were then dehydrated protected from light in completely filled 15 ml tubes under slow continuous rotation at room temperature through sequential ethanol / 2 % Tween-20 incubations: 30 % for four hours, 50 % for four hours, 70 % overnight, 90 % for four hours, absolute ethanol for four hours, and fresh absolute ethanol overnight. Dehydrated lungs were cleared in completely filled 5 ml amber polypropylene tubes containing MACS Clearing Solution for six hours at room temperature under slow continuous rotation, then transferred to MACS Clearing Solution, ethyl cinnamate (ECi), for imaging or stored at room temperature in polypropylene or glass tubes containing MACS Imaging Solution until imaging.

#### OptiMuS-prime tissue clearing (mouse brain)

Freshly dissected brains were collected in PBS buffer, fixed overnight at 4 °C in freshly prepared 4 % formaldehyde in PBS (ThermoFisher, 28908), and washed for 24 hours at room temperature with three PBS buffer changes; if not processed immediately, samples were stored in PBS/azide buffer at 4 °C. Fixed samples were passively cleared in OptiMuS-prime solution (100 mM Tris, 0.34 mM EDTA, 4 M urea, 10 % w/v D sorbitol, and 10 % w/v sodium cholate, pH 7.5, at 37 °C) under slow continuous rotation, using 15 ml solution per sample and daily solution exchange as needed until optically transparent. Samples were then incubated in permeabilisation solution (2 % Triton X-100/PBS) for 12 hours under slow continuous rotation and washed for one hour, followed by incubation in a blocking solution (10 % BSA/PBS) for 12 hours at 37 °C. The samples were immunostained with primary antibodies in staining solution (3 % (w/v) BSA and 2 % (v/v) Triton X-100 in PBS) for 5 days at 37 °C followed by washing for two hours in PBS and staining with secondary antibodies for four days at 37 °C, all under slow continuous rotation. The staining solution containing antibodies was replaced every 48 hours to maintain optimal labelling efficiency. Following staining, all samples were washed in PBS for six hours at room temperature under rotation and then immersed in OptiMuS RI-matching solution (100 mM Tris, 0.34 mM EDTA, 4 M urea, 10 % w/v D-sorbitol, and 75 % w/v Histodenz/iohexol, RI *≈* 1.47) for one day at 37 °C before imaging.

#### FOCM tissue clearing (mouse lung)

Mouse lung was first stained following the iDISCO immunostaining protocol. In short, samples were incubated first twice with 0.2 % Triton X-100 in PBS for two hours each time, following an overnight incubation in 20 % DMSO / 0.2 % Triton X-100 in PBS. The next day, the samples were incubated in 20 % DMSO / 0.2 % Triton X-100 / 0.2 % Tween 20 / 0.2 % sodium deoxycholate / 0.2 % NP40 in PBS. Samples were further washed in 0.2 % Triton X-100 in PBS twice for two hours. Permeabilisation was carried out by incubating the sample in 20 % DMSO / 0.2 % Triton X-100 / 0.3 M glycine in PBS overnight at 37 °C with shaking before incubating with Blocking Buffer (0.2 % Triton X-100 / 10 % DMSO / 6 % Normal Donkey Serum in PBS) for 24 hours at 37 °C with shaking. The samples were then incubated with primary antibodies (*α*-Smooth Muscle Actin, Abcam, ab124964, one in 200 dilution) diluted in primary staining buffer (5 % DMSO / 0.2 % Tween / 3 % Normal Donkey Serum / 10 µg/ml heparin in PBS) for four days. Lung was then washed four times at room temperature using 0.2 % Tween 20 with 10 µg/ml heparin in PBS (PTwH). Secondary antibodies (anti-rabbit AF568, Invitrogen, A10042) were then diluted in secondary staining buffer (0.2 % Tween / 3 % Normal Donkey Serum / 10 µg/ml heparin in PBS) and the sample was incubated for four more days at 37 °C. Finally, the sample was further washed in PTwH four times at room temperature for one hour each before clearing. FOCM clearing was performed according to the published protocol. In short, the stained lung was incubated with FOCM reagent (30 % urea (G Biosciences, BC89), 20 % d-sorbitol (ChemCruz, sc-203278), 5 % glycerol dissolved in DMSO) for three days at 37 °C with gentle shaking. After this period, the samples were transferred to an imaging dish for imaging.

#### Adipo tissue clearing (mouse small intestine)

Mouse gut tissues were collected from intracardially perfused animals and processed for whole mount staining and clearing following the modified AdipoClear tissue clearing and immunostaining protocol. Tissue was collected in 4 % PFA, left in fixation buffer overnight and transferred to PBS. Following three 30 min PBS washes, tissues were dehydrated at room temperature (RT) through a graded methanol (MeOH) series (20 %, 40 %, 60 %, 80 %, 100 %, and 100 %) prepared in B1n buffer (10 g glycine, 500 µl Triton X-100, 0.1 g sodium azide, and 500 ml Milli-Q water), with the 80 % MeOH step prepared in water. Samples were then delipidated overnight in a 2:1 dichloromethane (DCM):MeOH solution. Rehydration was performed sequentially in 100 % DCM, 100 % MeOH, 5:1 MeOH:H_2_O_2_ bleaching solution and decreasing concentrations of MeOH in B1n buffer before overnight incubation in 100 % B1n. All incubation steps were performed on a shaker. Tissue permeabilisation was carried out at room temprature using PTxwH buffer (500 µl Triton X-100, 250 µl Tween20, 0.1 g sodium azide, 1 mg heparin, 500 ml and PBS) supplemented with 5 % dimethyl sulfoxide (DMSO) and 0.3 M glycine, followed by washes in PTxwH buffer alone. Samples were incubated with primary antibodies diluted in PTxwH for 4–5 days at RT, washed extensively, and then incubated with secondary antibodies in PTxwH for 3–4 days at RT. Following immunostaining, tissues were dehydrated again through a graded MeOH series, delipidated overnight in 2:1 DCM:MeOH, washed in 100 % DCM, and cleared in ethyl cinnamate (ECI). Cleared gut tissues were stored in fresh ECI at room temperature in the dark until imaging.

#### Ce3D tissue clearing (mouse femur)

Mouse femurs were fixed for two hours with gentle shaking at 4 °C in freshly prepared 4 % formaldehyde in PBS (ThermoFisher, 28906). Samples were rinsed once with PBS and decalcified in 10 % EDTA (BDH, 205–358–3) (pH 7.8) at 4 °C for five days with gentle shaking. Samples were rinsed with PBS and processed for immunostaining and tissue clearing using the Ce3D kit (BioLegend, 427702) following the BioLegend protocol. Femurs were moved to a 24-well plate for permeabilisation and blocking using 1 ml of the Ce3D Permeabilisation/Blocking Buffer at RT with gentle shaking for two days. Samples were then transferred to a new well containing 1 ml primary antibody cocktail prepared in Ce3D Antibody Diluent Buffer and incubated at RT with gentle shaking for three days. Samples were washed three times in Ce3D wash buffer at RT with gentle shaking for eight hours. Samples were incubated with the secondary antibody cocktail prepared in Ce3D Antibody Diluent Buffer at RT with gentle shaking for three days. Samples were washed three times in Ce3D wash buffer at RT with gentle shaking for eight hours, with nuclear stain added during the second wash step. Samples were cleared using 1 ml Ce3D Tissue Clearing Solution for 24 hours at RT, transferred to a glass bottom plate with fresh Ce3D Tissue Clearing Solution, and immediately imaged.

#### REPLICA clearing (mouse kidney)

The full protocol for REPLICA clearing will be described in an upcoming manuscript by Cardoso Mestre et al.. After clearing, REPLICA samples were washed three times with PBS before being transferred into deionised water for long-term storage and imaging.

#### CUBIC clearing (mouse cerebellum)

CUBIC-R+ clearing was performed using the commercially available reagents CUBIC-L (T3740), CUBIC-R+ (T3741) and mounting media (M3294) sold by Tokyo Chemical Industry UK Ltd [21]. Following 4 % PFA fixation overnight at 4 °C, samples were washed three times with PBS for one hour each at room temperature. Prior to immunostaining, samples were delipidated by incubation with 50 % CUBIC-L at 37 °C overnight with gentle shaking, followed by incubation with 100 % CUBIC-L under the same conditions. This second incubation was performed for three days. Samples were then washed three times with PBS for two hours each at room temperature on a roller. Primary antibodies were diluted in staining buffer (3 % NDS / 0.25 % Triton X-100 in PBS) and samples were incubated at 37 °C for four days with gentle shaking. The primary staining solution was then removed and samples were washed three times with PBS for two hours each at room temperature. Secondary antibodies were diluted in staining buffer and incubated for a further four days at 37 °C with agitation. Samples were then washed three times with PBS for two hours each at room temperature. PBS was replaced with 1 % formaldehyde (diluted from 16 % stock; ThermoScientific, 28906) and samples were incubated overnight at 4 °C, then equilibrated to room temperature for one hour, followed by three PBS washes. Finally, refractive index was homogenised by incubating samples in 50 % CUBIC-R+ overnight at room temperature, followed by immersion in 100 % CUBIC-R+. This final incubation was performed for three days.

#### SmartBatch+ active tissue clearing and immunolabelling (mouse brain)

Samples were prepared for whole-tissue fluorescence imaging using a SHIELD-based tissue preservation, and SmartBatch+ Active tissue clearing protocol, that consisted of the next steps: delipidation, RADIANT primary and Standard secondary immunolabelling, and refractive-index-matching workflow adapted from LifeCanvas Technologies protocols. For SHIELD preservation, samples were transferred to freshly prepared SHIELD OFF solution consisting of 50 % SHIELD Epoxy Solution, 25 % SHIELD BufferSolution, and 25 % water by volume. Samples were incubated in SHIELD OFF solution at 4 °C with gentle shaking for 72 hours. Epoxy crosslinking was then performed by incubating samples in SHIELD ON Buffer for 24 hours at 37 °C with gentle shaking. Following SHIELD preservation, samples were stored in PBS containing 0.02 % sodium azide at 4 °C or proceeded directly to delipidation. For active delipidation, samples were first incubated in Delipidation Buffer for 24 hours at 45 °C with gentle shaking. If a 45 °C incubator was unavailable, samples were incubated at 37 °C. Samples were then mounted in mesh bags and loaded into a SmartBatch+ Clearing cup containing Delipidation Buffer. Conduction Buffer was added to the SmartBatch+ chamber, and clearing was performed using the Clearing Mode preset at 40 V, a 1250 mA current limit, and 42 °C. Clearing was performed for 30 hours. Following delipidation, samples were transferred to PBS containing 0.02 % sodium azide at 4 °C until immunolabelling. RADIANT primary immunolabelling was performed using a SmartBatch+ device equipped with the RADIANT upgrade. Samples were pre-incubated in SmartBatch+ RADIANT Buffer for 24 hours before labelling. Pre-incubation was performed either in 20 ml RADIANT Buffer in individual conical tubes, protected from light with gentle shaking at room temperature. On the day of labelling, the buffer was refreshed and samples were incubated for an additional one hour before loading into the device. For primary labelling, the medium-sized SmartBatch+ staining cup was selected according to the number of samples. The device chamber was filled with SmartBatch+ RADIANT Buffer according to cup size (85 ml for a medium cup). The staining cup was conditioned with RADIANT Buffer and then filled with 20 ml of RADIANT Buffer. FluoTag-X4 anti-RFP-AlexaFluor568 (NanoTag, 0404-AF568-L) was used for RADIANT immunolabelling of endogenous tdTomato expression in mouse brain tissue. A total of 100 µl of reconstituted FluoTag-X4 anti-RFP-AlexaFluor568 stock was added to 20 ml RADIANT staining solution. This corresponded to an approximate 1:200 dilution. The final total sdAb concentration was approximately 12.5 nM, with approximately 6.25 nM of each anti-RFP sdAb clone. The final dye concentration was approximately 25 nM. The estimated final mass concentration of the FluoTag conjugate was approximately µg/ml. This concentration was selected to approximate the molar binding-site concentration recommended for conventional anti-RFP IgG antibodies in the RADIANT protocol. The LifeCanvas protocol-recommended 18 µg IgG in 20 ml corresponds to approximately 12 nM antigen-binding sites, assuming a 150 kDa IgG with two binding arms. Because sdAb reagents are substantially smaller than full IgG antibodies, antibody concentration was compared by molarity rather than by mass concentration. Samples were placed in mesh bags, loaded into the staining cup, and fully submerged in the antibody-containing RADIANT Buffer. Labelling was performed using the Labeling 1 preset at 30 °C, 90 V, and a 350 mA current limit for 30 hours in a medium-size cup. Samples were protected from light throughout labelling and subsequent washes. After primary labelling, samples were transferred to PBS containing 0.02 % sodium azide and washed at room temperature with gentle shaking until the end of the day, with at least one buffer refresh. Samples were then fixed overnight at room teperature in conical tubes with 20 ml 4 % PFA in 1*×* PBS with gentleshaking, protected from light. For refractive-index matching, samples were equilibrated in EasyIndex solution with refractive index 1.52. EasyIndex was homogenised by gentle inversion before use. Samples were first incubated in 50 % EasyIndex and 50 % distilled water at 37 °C with gentle shaking in sealed, light-protected containers for 24 hours. Samples were then transferred to 100 % EasyIndex and incubated at 37 °C for the same duration or until optically transparent. Samples were maintained in 100 % EasyIndex until mounting and imaging, with latter performed in the fresh EasyIndex solution.

