## Supplementary material for "Index-agnostic oblique plane light sheet microscopy of centimetre-scale cleared tissues at subcellular resolution"

### A Calculation of the wavefront for a displaced object point

Following the derivation of Botcherby et al., the origin of our coordinate system is set to the on-axis focal point of an objective lens [1]. We now consider a point on the principal sphere given by $\boldsymbol{f}$ and look at the optical path length difference, $\Delta$, obtained when reaching this point from a displaced object point, $\boldsymbol{r}$, for which the vector to reach the same principal sphere point is $\boldsymbol{w}$. Hence, we have:

$$\Delta = |\boldsymbol{w}| - |\boldsymbol{f}| = |\boldsymbol{f} - \boldsymbol{r}| - f = \sqrt{f^2 - 2\boldsymbol{f}\cdot\boldsymbol{r} + r^2} - f. \tag{S1}$$

Using spherical polar coordinates, we then have:

$$\Delta = f\sqrt{1 - \frac{2}{f}\left(x\sin\theta\cos\phi + y\sin\theta\sin\phi + z\cos\theta\right) + \frac{r^2}{f^2}} - f, \tag{S2}$$

where $\theta$ is the polar angle and $\phi$ the azimuthal angle.

In the standard analysis of remote refocussing, this is then expanded to first order under the assumption that $r \ll f$ [1]. Introducing the sine condition ($\sin\theta = \rho\sin\alpha$), multiplying by the vacuum wavenumber, $k$, and including the possibility that the object point is immersed in a medium of refractive index, $n$, we obtain the phase profile in the pupil plane:

$$\Psi_{\text{approx}}\left(\rho,\phi;\boldsymbol{r}\right) \approx nk\sin\alpha\left(x\rho\cos\phi + y\rho\sin\phi + z\sqrt{\frac{1}{\sin^2\alpha} - \rho^2}\right), \tag{S3}$$

where $\phi$ is now the azimuthal angle in the pupil rather than in the object space. This pupil function is odd, i.e. $\Psi_{\text{rr}}\left(\rho,\phi;\boldsymbol{r}\right) = -\Psi_{\text{rr}}\left(\rho,\phi;-\boldsymbol{r}\right)$ and so it would be possible to image all points in a 3D volume stigmatically. However, as noted in the original remote refocussing publications, higher order terms ignored by this first-order approximation do not cancel out and so the real pupil function does not have this odd parity. While an expansion to second order was used to analyse the volume over which a high Strehl ratio is maintained in the original remote refocussing publications, we instead chose to investigate the full pupil function on-axis (i.e. $x = y = 0$):

$$\Psi_{\text{rr}}\left(\rho;z\right) = nkf\left(\sqrt{1 - \frac{2z}{f}\sqrt{1-\rho^2\sin^2\alpha} + \frac{z^2}{f^2}} - 1\right). \tag{S4}$$

Imaging through a depth, $d$, into a refractive index, $n_{\text{sample}}$, different to the one the lens was designed for, $n_{\text{design}}$, across a boundary normal to the optical axis, leads to a further pupil phase function [2]:

$$\Psi_{\text{mismatch}}\left(\rho;d\right) = kn_{\text{design}}d\left(\sqrt{\frac{n_{\text{sample}}^2}{n_{\text{design}}^2} - \rho^2\sin^2\alpha} - \sqrt{1-\rho^2\sin^2\alpha}\right). \tag{S5}$$

Altogether, the overall pupil phase function is then given by:

$$\Psi\left(\rho;z,d\right) = n_{\text{design}}k\left[f\left(\sqrt{1 - \frac{2z}{f}\sqrt{1-\rho^2\sin^2\alpha} + \frac{z^2}{f^2}} - 1\right) + d\left(\sqrt{\frac{n_{\text{sample}}^2}{n_{\text{design}}^2} - \rho^2\sin^2\alpha} - \sqrt{1-\rho^2\sin^2\alpha}\right)\right]. \tag{S6}$$

We now assume that this pupil is imaged, via a $4f$ relay, into the pupil of a second objective lens operating in a uniform refractive index, $n_{\text{remote}}$. For this lens, the pupil function for a point along the optical axis displaced by $z_2$ from the focal plane would then be:

$$\Psi_2\left(\rho;z_2\right) = n_{\text{remote}}kf_2\left(\sqrt{1 - \frac{2z_2}{f_2}\sqrt{1-\rho^2\sin^2\alpha} + \frac{z_2^2}{f_2^2}} - 1\right). \tag{S7}$$

For an overall lateral system magnification $M$, we have an axial magnification of $M^2 n_{\text{remote}}/n_{\text{sample}}$. Hence, the wavefront produced by the first objective by a point displaced by $z$ should be compared with the wavefront produced by the second objective for a point displaced by $zM^2 n_{\text{remote}}/n_{\text{sample}}$, and so we wish to evaluate:

$$\begin{aligned}\Psi_{\text{overall}}\left(\rho;z,d\right) =& n_{\text{design}}k\left[f\left(\sqrt{1 - \frac{2z}{f}\sqrt{1-\rho^2\sin^2\alpha} + \frac{z^2}{f^2}} - 1\right) + d\left(\sqrt{\frac{n_{\text{sample}}^2}{n_{\text{design}}^2} - \rho^2\sin^2\alpha} - \sqrt{1-\rho^2\sin^2\alpha}\right)\right] \\ &- n_{\text{remote}}kf_2\left(\sqrt{1 - \frac{2zM^2 n_{\text{remote}}/n_{\text{sample}}}{f_2}\sqrt{1-\rho^2\sin^2\alpha} + \frac{z^2M^4 n_{\text{remote}}^2/n_{\text{sample}}^2}{f_2^2}} - 1\right).\end{aligned} \tag{S8}$$

However, we must first remove any piston:

$$\Psi_p = 1, \tag{S9}$$

and pistonless defocus:

$$\Psi_d = n_{\text{design}} k \sqrt{1-\rho^2 \sin^2\alpha} - \frac{2 n_{\text{design}} k}{3\sin^2\alpha}\left(1-\cos^3\alpha\right). \tag{S10}$$

The amounts of these functions contained within the wavefront can be found via calculation of the inner product,

$$\delta = \frac{\int \Psi_{\text{overall}} \Psi_i \rho \,\mathrm{d}\rho}{\int \Psi_i \Psi_i \rho \,\mathrm{d}\rho}. \tag{S11}$$

For piston, we obtain a coefficient of:

$$\begin{aligned}\delta_p = \frac{1}{15} k n_{\text{design}} \Bigg( &-\frac{2\csc^2(\alpha)\left(\left(f^2+3fz+z^2\right)|f-z|^3 - \sqrt{f^2-2fz\cos(\alpha)+z^2}\left(f^4+fz\left(\cos(\alpha)\left(f^2+z^2\right)-3fz\cos(2\alpha)\right)-f^2z^2+z^4\right)\right)}{f^2z^2} \\ &- 10d\left(\cos(\alpha)+\tan\left(\frac{\alpha}{2}\right)\csc(\alpha)\right) + \frac{10d\csc^2(\alpha)\left(n_{\text{sample}}^3 - \left(n_{\text{sample}}^2 - n_{\text{design}}^2\sin^2(\alpha)\right)^{3/2}\right)}{n_{\text{design}}^3} - 15f \Bigg),\end{aligned} \tag{S12}$$

which, as expected, does not depend on $f_2$, $M$ or $n_{\text{remote}}$, and evaluates to $k f n_{\text{design}}$ when $z = d = 0$.

A more protracted calculation, best accompanied by the consumption of multiple cups of tea, leads to the following coefficient for defocus:

$$\begin{aligned}\delta_d = \frac{\csc^2\alpha}{420 f^3 n_{\text{design}}^4 z^3\,(3+\cos 2\alpha)} \Bigg[ &-35 f^3 n_{\text{design}}\left(32 f n_{\text{design}}^3 + 3d\left(5n_{\text{design}}^3 - 4n_{\text{design}}^2 n_{\text{sample}} - 4n_{\text{sample}}^3\right)\right) z^3 \\ &- 32 n_{\text{design}}^4\left(2f^4+6f^3z+19f^2z^2+6fz^3+2z^4\right)|f-z|^3 \\ &+ 1120 f^4 n_{\text{design}}^4 z^3\cos^3\alpha \\ &+ 16 n_{\text{design}}^4\left(f^2+z^2\right)\left(4f^4+11f^2z^2+4z^4\right)\sqrt{f^2+z^2-2fz\cos\alpha} \\ &+ 2 f n_{\text{design}} z\cos\alpha\Bigg(16 n_{\text{design}}^3\left(2f^4-11f^2z^2+2z^4\right)\sqrt{f^2+z^2-2fz\cos\alpha} \\ &\quad - 15 f^2 z^2\Bigg(7 d n_{\text{sample}}^2\sqrt{-2n_{\text{design}}^2+4n_{\text{sample}}^2+2n_{\text{design}}^2\cos 2\alpha} \\ &\quad\quad + n_{\text{design}}^2\cos 2\alpha\left(16 n_{\text{design}}\sqrt{f^2+z^2-2fz\cos\alpha} + 7d\sqrt{-2n_{\text{design}}^2+4n_{\text{sample}}^2+2n_{\text{design}}^2\cos 2\alpha}\right)\Bigg)\Bigg) \\ &+ 3f^2z^2\Bigg(4n_{\text{design}}^4\left(35dfz+4\left(f^2+z^2\right)\sqrt{f^2+z^2-2fz\cos\alpha}\right)\cos 2\alpha \\ &\quad + 35dfz\left(n_{\text{design}}^4\cos 4\alpha - 4\left(n_{\text{design}}^2-n_{\text{sample}}^2\right)^2\log\left(\frac{n_{\text{design}}+n_{\text{sample}}}{n_{\text{design}}\cos\alpha+\sqrt{n_{\text{sample}}^2-n_{\text{design}}^2\sin^2\alpha}}\right)\right)\Bigg)\Bigg],\end{aligned} \tag{S13}$$

which again, as expected, does not depend on $f_2$, $M$ or $n_{\text{remote}}$.

As these equations are somewhat cumbersome, in practice we instead calculated the appropriate coefficients numerically. Nevertheless, we felt we should share the results with the community to prevent any future researchers embarking on the same foolhardy journey.

Overall, our wavefunction of interest is given by:

$$\Psi_{\text{overall}}(\rho; z, d) = n_{\text{design}} k \left[ f\left(\sqrt{1 - \frac{2z}{f}\sqrt{1-\rho^2\sin^2\alpha} + \frac{z^2}{f^2}} - 1\right) + d\left(\sqrt{\frac{n_{\text{sample}}^2}{n_{\text{design}}^2} - \rho^2\sin^2\alpha} - \sqrt{1-\rho^2\sin^2\alpha}\right)\right]$$
$$- n_{\text{remote}} k f_2 \left(\sqrt{1 - \frac{2zM^2 n_{\text{remote}}/n_{\text{design}}}{f_2}\sqrt{1-\rho^2\sin^2\alpha} + \frac{z^2M^4 n_{\text{remote}}^2/n_{\text{design}}^2}{f_2^2}} - 1\right)$$
$$- \delta_p - \delta_d \left[ n_{\text{design}} k \sqrt{1-\rho^2\sin^2\alpha} - \frac{2n_{\text{design}}k}{3\sin^2\alpha}\left(1-\cos^3\alpha\right)\right], \quad \text{(S14)}$$

and we wish to calculate the Strehl ratio:

$$S = \left|\frac{1}{\pi}\int \exp\left(i\Psi_{\text{overall}}\right)\rho\,\mathrm{d}\rho\,\mathrm{d}\phi\right|^2. \quad \text{(S15)}$$

#### B Sheet tilt as a function of refractive index

In traditional oblique plane microscopes the lateral and axial magnifications of the remote refocussing relay is equal to the sample refractive index. Therefore, the sheet angle in the sample is conserved in the remote space. However, in CtDvOPM the lateral magnification remains fixed despite imaging samples with refractive indices ranging from 1.33 to 1.56. This leads to a sample refractive-index-dependent axial magnification and hence axial stretching, which causes the effective imaging angle in the sample to change. Therefore, we need to calculate the exact imaging angle for each refractive index so that we can rotate the light sheet launch to the correct angle such that the excitation plane is coplanar with the camera detection plane.

For an imaging system with a lateral magnification, $M_{\text{lat}}$, the axial magnification, $M_{\text{ax}}$, is given by:

$$M_{\text{ax}} = \frac{n_{\text{sample}}}{n_{\text{image}}} M_{\text{lat}}^2, \quad \text{(S16)}$$

where, $n_{\text{sample}}$ and $n_{\text{image}}$ are the refractive indices in the sample and imaging space, respectively. As shown in Supplementary Figure 1, if we consider a distance along the $x$ and $z$ axes in the sample space of $\Delta x$ and $\Delta z$ the imaging angle in the sample space, $\theta$, is given by,

$$\theta = \arctan\left(\frac{\Delta z}{\Delta x}\right). \quad \text{(S17)}$$

Similarly, the matched plane in the imaging space has the angle, $\theta'$, given by:

$$\theta' = \arctan\left(\frac{\Delta z'}{\Delta x'}\right). \quad \text{(S18)}$$

By using equations S16 and S17, and assuming that $n_{\text{image}} = 1$, we can rewrite equation S17 as a function of the camera angle, $\theta'$, sample refractive index, $n_{\text{sample}}$, and the optical system's magnification, $M_{\text{lat}}$, such that:

$$\theta = \arctan\left(\frac{n_{\text{sample}}}{M_{\text{lat}}}\tan\theta'\right). \quad \text{(S19)}$$

Using Snell's law, the angle of incidence required for the laser launch, $\phi$, assuming a dry immersion lens, is given by:

$$\phi = \arcsin\left[n_{\text{sample}}\sin\left(\frac{\pi}{2} - \arctan\left(\frac{n_{\text{sample}}}{M_{\text{lat}}}\tan\theta'\right)\right)\right]. \quad \text{(S20)}$$

The equations S19 and S20 are plotted over the range 1.00–2.00 for $n_{\text{sample}}$, for a lateral magnification and camera angle corresponding to our implementation (1.45, 50°) in Supplementary Figure 2. Over this entire range, the change in tilt is less than 18°.

#### C Sheet aberration as a function of refractive index

In CtDvOPM, the light sheet is generated using a dry objective inclined relative to the coverglass. Consequently, the excitation beam crosses a refractive-index boundary at a large incidence angle before forming the light sheet in the sample. Although the numerical aperture of the excitation light sheet is relatively low, the large incidence angle at the coverglass can still introduce significant aberrations into the excitation field. To quantify these aberrations, we first consider the pupil phase function associated with focussing through a refractive-index mismatch. We then transform this phase profile into the coordinate frame of the excitation objective and evaluate it over the excitation pupil.

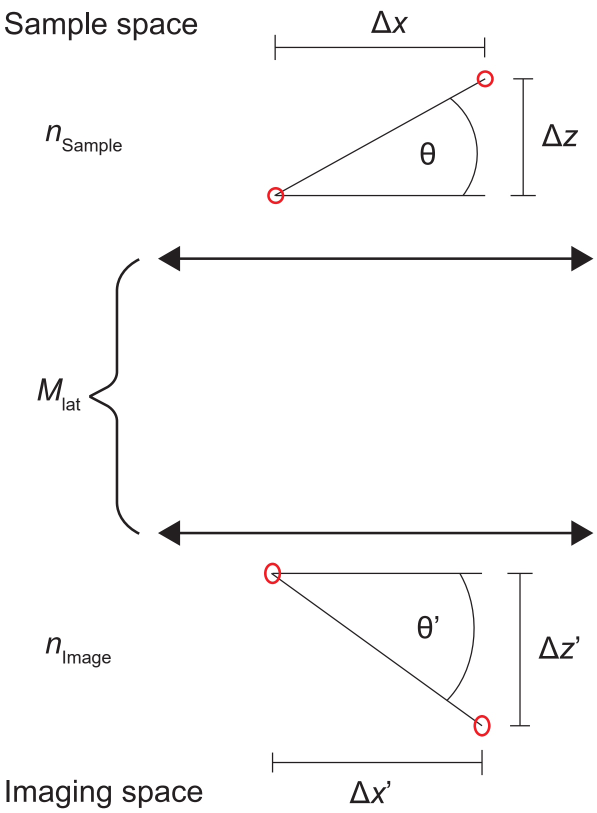


**Supplementary Figure 1: Diagram illustrating the mapping of planes in CtDvOPM.**
An optical system with the lateral magnification $M_{lat}$ images the two red points in a refractive index of $n_1$ separated by $\Delta x$ and $\Delta y$ in to an image space of refractive index $n_2$. In the imaging space the two red points are separated by $\Delta x'$ and $\Delta y'$. When the lateral magnification equals $\frac{n_1}{n_2}$ all angles are conserved, otherwise there is an axial stretching, effectively rotating the imaging angle in the sample.

Focussing into a medium of refractive index $n_{\text{sample}}$ using an objective designed for refractive index $n_{\text{design}}$, through a depth $d$ across a refractive-index boundary normal to the optical axis, produces the pupil phase function:

$$\Psi_{\text{mismatch}}(\rho; d) = kn_{\text{design}} d \left( \sqrt{\frac{n^2_{\text{sample}}}{n^2_{\text{design}}} - \rho^2 \sin^2\alpha} - \sqrt{1 - \rho^2 \sin^2\alpha} \right). \tag{S21}$$

Here, $\alpha$ does not correspond to the angular aperture of the excitation objective. Instead, it represents the maximum angle subtended by rays in the excitation cone relative to the surface normal. This angle is given by the sum of the incidence angle of the excitation sheet chief ray at the coverglass, $\phi$, and the angular aperture of the excitation objective, $\alpha_{\text{ex}}$,

$$\begin{aligned} \alpha &= \phi + \alpha_{\text{ex}}, \\ &= \arcsin\left[\frac{n_{\text{sample}} \sin\left(\frac{\pi}{2} - \theta\right)}{n_{\text{design}}}\right] + \arcsin\left[\frac{\text{NA}}{n_{\text{design}}}\right], \\ &= \arcsin\left[\frac{n_{\text{sample}} \sin\left(\frac{\pi}{2} - \arctan\left(\frac{n_{\text{sample}}}{M_{\text{lat}}} \tan\theta'\right)\right)}{n_{\text{design}}}\right] + \arcsin\left[\frac{\text{NA}}{n_{\text{design}}}\right]. \end{aligned} \tag{S22}$$

where $\theta$ is angle of the sheet in the sample, given by S19 and NA is the numerical aperture of the sheet in the design refractive index, $n_{\text{design}}$. To express the mismatch phase function in the coordinate system of the tilted excitation pupil, we map the pupil coordinates onto the reference Ewald sphere and rotate this sphere into the excitation frame. We first perform an orthographic projection of the original pupil coordinates $(x,y)$ onto a sphere of radius $R = n_{\text{design}} / \sin\alpha$ such that the pupil coordinates $(x,y)$ are mapped onto the reference sphere coordinates $(x', y', z')$ according to

$$\begin{aligned} x' &= x, \\ y' &= y, \\ z' &= \sqrt{R^2 - x^2 - y^2}. \end{aligned} \tag{S23}$$

The reference sphere is then rotated about the y-axis by the excitation incidence angle $\phi$, giving transformed coordinates

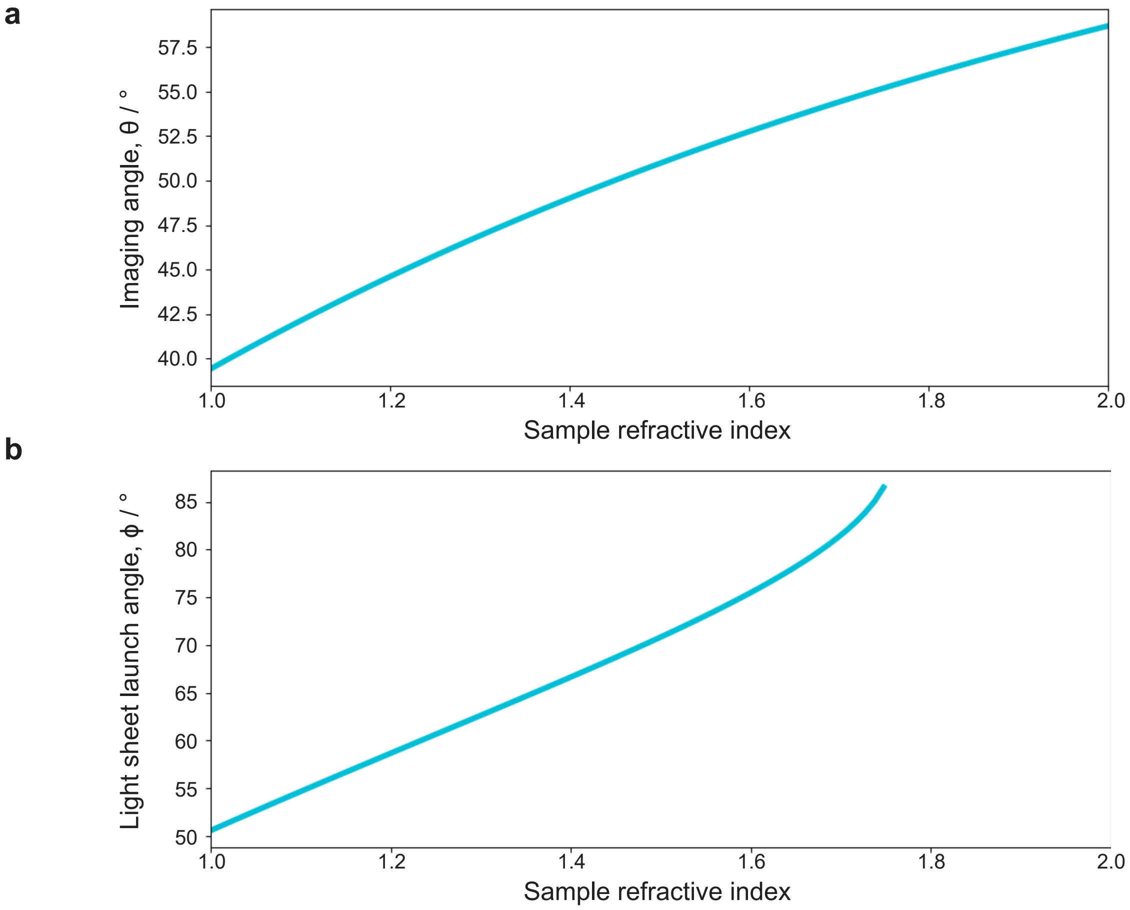


**Supplementary Figure 2: Imaging angle as a function of refractive index for CtDvOPM.**
**a** Plot showing the imaging angle as a function of sample refractive index for an imaging system with a lateral magnification of 1.45 and a camera angle of 50°. **b** Corresponding plot of incident light sheet angle for imaging samples over the refractive index range shown in a.

$(X, Y, Z)$:

$$\begin{bmatrix} X \\ Y \\ Z \end{bmatrix} = \begin{bmatrix} \cos\phi & 0 & \sin\phi \\ 0 & 1 & 0 \\ -\sin\phi & 0 & \cos\phi \end{bmatrix} \begin{bmatrix} x \\ y \\ \sqrt{R^2 - x^2 - y^2} \end{bmatrix} = \begin{bmatrix} x\cos\phi + \sin\phi\sqrt{R^2 - x^2 - y^2} \\ y \\ \cos\phi\sqrt{R^2 - x^2 - y^2} - x\sin\phi \end{bmatrix}. \tag{S24}$$

Performing an orthographic projection from the rotated sphere onto the excitation pupil plane therefore gives the coordinate transformation between the original pupil coordinates $(x, y)$ and the excitation pupil coordinates $(X, Y)$:

$$\begin{aligned} X &= x\cos\phi + \sin\phi\sqrt{R^2 - x^2 - y^2}, \\ Y &= y. \end{aligned} \tag{S25}$$

Rearranging these expressions gives the inverse transformation from the excitation pupil coordinates back to the original pupil coordinates,

$$\begin{aligned} x &= X\cos\phi - \sin\phi\sqrt{R^2 - X^2 - Y^2}, \\ y &= Y. \end{aligned} \tag{S26}$$

Substituting this transformation into equation S21, and assuming cylindrical focussing such that the system is invariant along the $y$-direction ($y = Y = 0$), gives the excitation sheet wavefront

$$\Psi_{\text{sheet}}(X; d) = kn_{\text{design}}d\left(\sqrt{\frac{n^2_{\text{sample}}}{n^2_{\text{design}}} - \left(X\cos\phi - \sin\phi\sqrt{R^2 - X^2}\right)^2 \sin^2\alpha} - \sqrt{1 - \left(X\cos\phi - \sin\phi\sqrt{R^2 - X^2}\right)^2 \sin^2\alpha}\right) \tag{S27}$$

The coordinate $X$ is evaluated over the interval

$$-\frac{n_{\text{design}}\sin\alpha_{\text{ex}}}{\sin\alpha} \leq X \leq \frac{n_{\text{design}}\sin\alpha_{\text{ex}}}{\sin\alpha}. \tag{S28}$$

Before calculating the Strehl ratio, we need to remove the contributions of the wavefront that do not affect the shape of the PSF, namely piston, tip and defocus. For piston, we have

$$\Psi_p = 1, \tag{S29}$$

while for tip:

$$\Psi_t = X, \tag{S30}$$

and for pistonless defocus:

$$\Psi_d = n_{\text{design}}k\sqrt{1 - X^2\sin^2\alpha_{\text{ex}}} - \frac{2n_{\text{design}}k}{3\sin^2\alpha_{\text{ex}}}\left(1 - \cos^3\alpha_{\text{ex}}\right). \tag{S31}$$

This gives a residual wavefront that can be used to calculate the Strehl ratio:

$$\Psi_{\text{residual}} = \Psi_{\text{sheet}} - \delta_p - \delta_t\Psi_t - \delta_d\Psi_d. \tag{S32}$$

where the values $\delta_p$, $\delta_t$ and $\delta_d$ are the coefficients for piston, tip and defocus respectively. For the 1D cyclindrical lens case, the Strehl ratio is given by:

$$S = \left|\frac{\sin\alpha}{2n_{\text{design}}\sin\alpha_{\text{ex}}}\int_{-n_{\text{design}}\sin\alpha_{\text{ex}}/\sin\alpha}^{n_{\text{design}}\sin\alpha_{\text{ex}}/\sin\alpha} \exp\left(i\Psi_{\text{sheet}}(x)\right)\mathrm{d}x\right|^2. \tag{S33}$$

Here, we solve for the aberration coefficients numerically by performing a least-squares linear regression to independently fit equations S29–S31 to the excitation-sheet wavefront, $\Psi_{\text{sheet}}$. This approach is possible because the piston, tip, and defocus functions are orthogonal. Using equations S32 and S33, we then evaluate the performance of the excitation sheet under different imaging conditions. Supplementary Figure 3 presents the predicted sheet performance when focussing 1 mm along $z$ into samples with refractive indices of 1.33, 1.45, and 1.56, using excitation-sheet NAs in air of up to 0.1. The results show that, at this depth, a sheet NA of 0.06 in air, as used throughout this work, remains diffraction limited across all sample refractive indices considered.

However, when imaging into water ($n = 1.33$), using the full illumination NA of 0.1 causes the excitation sheet to no longer remain diffraction limited. Interestingly, the excitation sheet remains diffraction limited at higher input NAs when imaging into higher-refractive-index samples. At first glance, this appears counterintuitive, since imaging across a larger refractive-index mismatch would normally be expected to increase aberrations.

In this case, however, the opposite behaviour occurs because the sheet intersects the refractive-index boundary at a large angle. Refraction at the interface compresses the angular extent of the excitation cone within the sample, thereby reducing the resulting wavefront aberration. This behaviour is also evident in figure Supplementary Figure 3b. Although the Strehl ratio indicates that the focus remains diffraction limited, the full width at half maximum (FWHM) is significantly larger than would be expected from the input NA in air. This is because the inclined excitation cone effectively reduces the NA of the sheet within the sample. Therefore, the sheet is not considered aberrated but has become broadened. Consequently, the effective sheet NA in the sample differs from the input NA in air, and this effect must therefore be taken into account when selecting the illumination conditions.

Axially swept light-sheet microscopy (ASLM) is often used to achieve increased axial resolution over extended imaging volumes through the use of a high-NA excitation sheet. Although CtDvOPM is theoretically compatible with ASLM, the results of the present analysis indicate that, when launching the excitation beam directly from air into the sample, adaptive optics would likely be required to correct the aberrations introduced at higher illumination NAs, as the required wavefront correction is a function of depth within the sample. In particular, the asymmetric ringing pattern observed in figure Supplementary Figure 3c suggests that coma is the dominant aberration under these conditions. To verify this, we removed primary coma from the residual phase profile and noted in our simulations that this alone was sufficient render the light sheet diffraction limited for all the imaging conditions shown in Supplementary Figure 3.

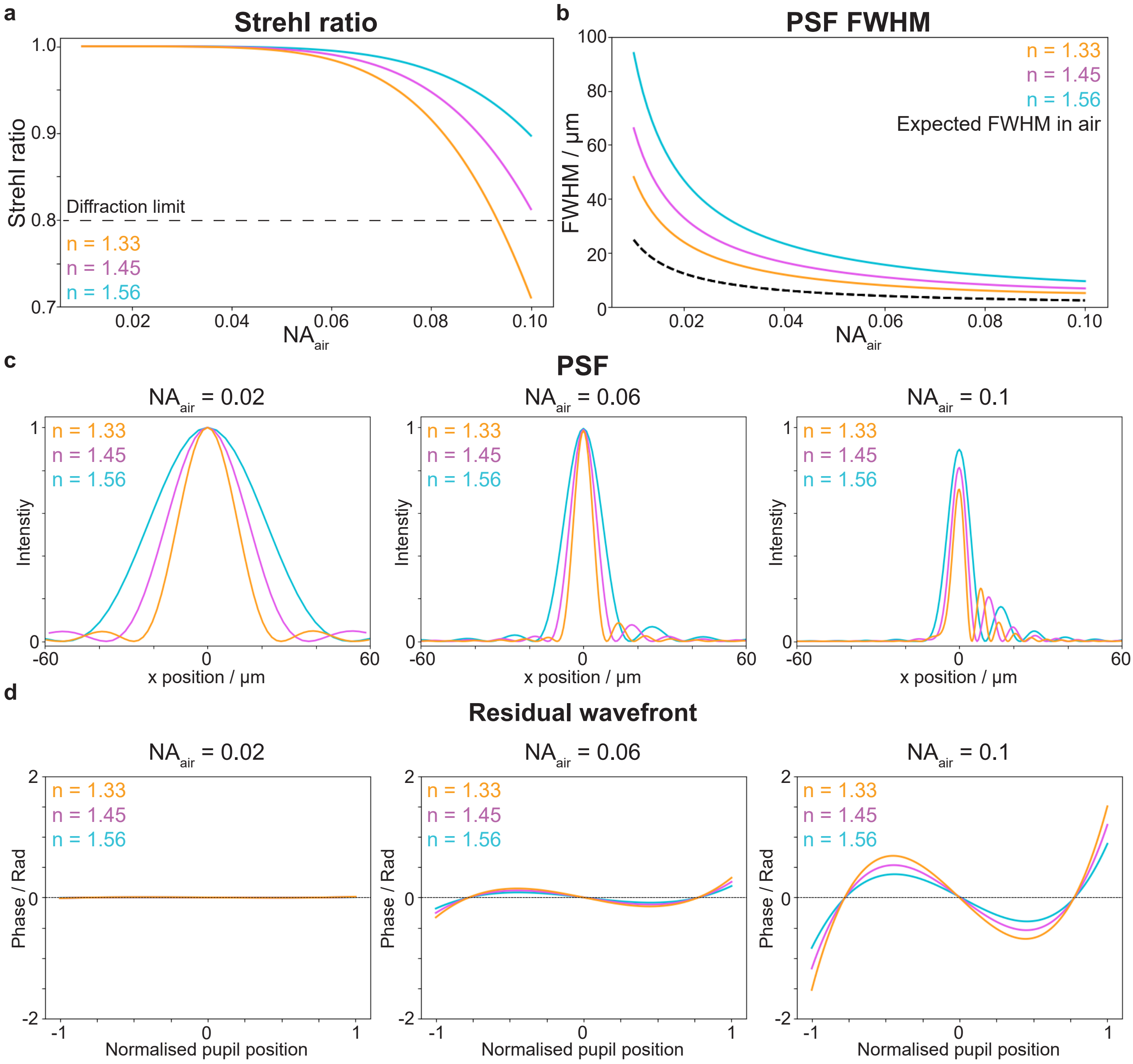


**Supplementary Figure 3: Computational analysis of lightsheet performance when imaging into different refractive indices (n = 1.33–1.56).**
**a** A plot of the Strehl ratio of a cylindrical focus at a $z$ depth of 1 mm when imaging into n = 1.33 (yellow), n = 1.45 (magenta) and n = 1.56 (cyan) for input NAs in air from 0.01 – 0.1. **b** A plot showing the FWHM under the same imaging conditions. The black dotted line represents the expected FWHM based on the NA in air. **c** Intensity cross sections of the PSF when the input NA is 0.02, 0.06 and 0.1 respectively. **d** Residual wavefront plots of the PSFs shown in **c**.

### D Parts list for CtDvOPM

| Supplier | Part no. | Description | QTY | Price / € | Subtotal / € |
|---|---|---|---|---|---|
| | | *Optics* | | | |
| Edmund Optics | 15-872 | 0.69× CobaltTL Telecentric Lens | 1 | 2820.00 | 2820.00 |
| Thorlabs | RC12APC-P01 | Reflective Collimator, RFL = 50.8 mm | 1 | 1093.01 | 1093.01 |
| Thorlabs | TL2X-SAP | 2× Objective, 0.1 NA, 56.3 mm WD | 1 | 1380.24 | 1380.24 |
| Thorlabs | ACY254-100-A | $f$ = 100 mm, Cylindrical Achromat | 1 | 434.44 | 434.44 |
| Thorlabs | BB1-E02 | 1-inch Broadband Dielectric Mirror | 2 | 79.00 | 158.00 |
| Thorlabs | WG41010-A | 1-inch UVFS Broadband Window, t = 1 mm | 1 | 100.87 | 100.87 |
| | | *Optomechanics* | | | |
| Thorlabs | KCB1C/M | 30 mm right-angle kinematic mirror mount | 2 | 87.24 | 174.48 |
| Thorlabs | CP33/M | 30 mm cage plate | 4 | 18.57 | 74.28 |
| Thorlabs | CP36 | 30 mm double-bore cage plate | 2 | 22.89 | 45.78 |
| Thorlabs | CBB1/M | 30 mm cage system u-bench | 1 | 87.24 | 87.24 |
| Thorlabs | ER6 | 6-inch long cage rod | 4 | 9.11 | 36.44 |
| Thorlabs | ER3 | 3-inch long cage rod | 4 | 6.94 | 27.76 |
| Thorlabs | ER2 | 2-inch long cage rod | 4 | 6.41 | 22.64 |
| Thorlabs | RS75/M | 25 mm pillar post, L = 75 mm | 4 | 29.12 | 116.48 |
| Thorlabs | RS150/M | 25 mm pillar post, L = 150 mm | 4 | 30.71 | 146.84 |
| Thorlabs | P200/M | 1 inch Mounting Post, M6 Taps, L = 200 mm | 4 | 65.12 | 260.48 |
| Thorlabs | TR20/M | 12.7 mm optical post, L = 20 mm | 4 | 5.24 | 20.96 |
| Thorlabs | PH20/M | 12.7 mm post holder, L = 20 mm | 4 | 7.94 | 31.76 |
| Thorlabs | SM1A12 | External SM1 and Internal M25 × 0.75 adapter | 1 | 23.34 | 23.34 |
| Thorlabs | SM1L30 | 3-inch SM1 lens tube | 1 | 28.43 | 28.43 |
| Thorlabs | SM1L05 | 0.5-inch SM1 lens tube | 1 | 13.24 | 13.24 |
| Thorlabs | AB90H | Slim right-angle bracket | 1 | 29.74 | 29.74 |
| Thorlabs | AP90RL/M | Large right-angle bracket | 1 | 189.50 | 189.50 |
| Thorlabs | B3060A | Nexus breadboard | 1 | 794.83 | 794.83 |
| Thorlabs | CRM1PT/M | Precision cage rotation mount | 1 | 233.39 | 233.39 |
| Thorlabs | AV4/M | Sorbothane Feet 4-pieces | 1 | 26.00 | 26.00 |
| | | *Translation stages* | | | |
| Thorlabs | XR25P/M | 25 mm linear translation stage | 1 | 488.91 | 488.91 |
| Thorlabs | XRN25C/M | Compact 25 mm linear translation stage | 1 | 456.49 | 456.49 |
| Thorlabs | MVS1/M | 25 mm vertical translation stage | 1 | 955.63 | 955.63 |
| Thorlabs | LX20/M | 25 mm XY translation stage | 1 | 962.29 | 962.29 |
| Thorlabs | ZFS25B | 25 mm travel stepper motor actuator | 2 | 1335.65 | 2671.30 |
| Thorlabs | KDC101 | K-Cube brushed DC servo motor controller | 2 | 751.52 | 1503.04 |
| | | *Camera* | | | |
| IDS | U3-31R2SE-M | Monochrome camera, IMX540 sensor | 1 | 2540.35 | 2540.35 |
| Eureca | N/A | Camera coverglass removal | 1 | 480.00 | 480.00 |
| | | *Beam shifter* | | | |
| Edmund Optics | 23-851 | 20mm Optotune Beam Shifter | 1 | 840.00 | 840.00 |
| Edmund Optics | 23-717 | Optotune Current Controller | 1 | 1912.38 | 1912.38 |
| Edmund Optics | 22-410 | Optotune ICC-4C Power Adapter | 1 | 69.00 | 69.00 |
| Edmund Optics | 23-718 | Optotune ICC-4C-2000 Extension Kit | 1 | 231.00 | 231.00 |
| | | *Laser* | | | |
| Cobolt | Skyra | 4-channel fibre coupled laser | 1 | 17334.47 | 17334.47 |
| | | *Filter wheel* | | | |
| Cairn Research | P1025/006/000 | OptoSpin six-position filter wheel | 1 | 1594.26 | 1594.26 |
| Cairn Research | P1020/006/000 | OptoSpin controller | 1 | 1912.38 | 1912.38 |
| | | *Custom parts — quote for reference only* | | | |
| Xometry | N/A | CNC Machined parts | 1 | 1779.67 | 1779.67 |
| Xometry | N/A | 3D printed parts | 1 | 35.56 | 35.56 |
| | | **Total cost** | | | **44403.92** |

### E Deskewing raw data into the laboratory frame

Raw data is acquired in a coordinate frame with respect to the angled focal plane in the sample $(x', y', z')$, as defined in Supplementary Figure 4. Therefore, to convert the raw data volume into a stack in the lab coordinates $(x,y,z)$ the data must

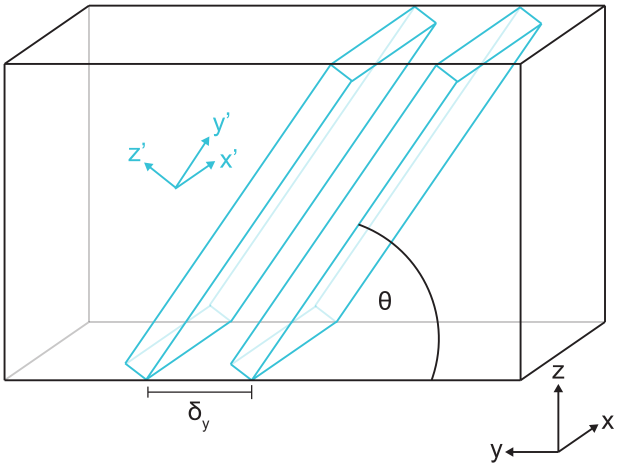


**Supplementary Figure 4: Geometry for data acquisition in CtDvOPM.**
CtDvOPM data are acquired in the rotated coordinate frame $(x', y', z')$, shown in blue. Raw frames are rotated by an angle $\theta$ about the $x$ axis and translated by a distance $\delta_y$ along the $y$ axis, where $(x, y, z)$ denote the laboratory coordinate system. The raw data are subsequently transformed into the laboratory frame through deskewing.

be deskewed. Whilst this deskewing process is the same as for other light sheet systems, the CtDvOPM has a sample refractive index-dependent compression of the imaging angle that must be accounted for.

When imaging into a refractive index that is less (more) than the magnification of the system the imaging depth is compressed (elongated), resulting in an anisotropic voxel size. To correct this, the raw data is first stretched along the $y'$ direction using,

$$\begin{bmatrix} 1 & 0 & 0 \\ 0 & S_y & 0 \\ 0 & 0 & 1 \end{bmatrix} \begin{bmatrix} x' \\ y' \\ z' \end{bmatrix} \tag{S34}$$

where, the stretch factor, $S_y$, is given by,

$$S_y = \frac{\cos\theta'}{\cos\theta} \tag{S35}$$

where $\theta$ and $\theta'$ are the angles of the focal plane in the sample and remote space respectively. Following this, the dataset is sheared along the $y'$ axis to account for the lateral shift along this axis induced when the sample is scanned along the $y$ axis during acquisition. This is achieved with the matrix,

$$\begin{bmatrix} 1 & 0 & 0 \\ 0 & 1 & \gamma \\ 0 & 0 & 1 \end{bmatrix} \tag{S36}$$

where, the shear factor, $\gamma$, is given by,

$$\gamma = -\frac{M_{\mathrm{lat}}\delta_y \cos\theta}{p_x} \tag{S37}$$

where $\delta_y$ is the step size along the $y$ axis during acquisition, $p_x$ is the physical pixel size and $M_{\mathrm{lat}}$ is the lateral magnification of the imaging system.

To make the voxels isotropic the dataset is then scaled along the $z'$ axis using,

$$\begin{bmatrix} 1 & 0 & 0 \\ 0 & 1 & 0 \\ 0 & 0 & \epsilon \end{bmatrix} \tag{S38}$$

where, the scale factor, $\epsilon$, is equal to

$$\epsilon = \frac{\delta_y \sin\theta M_{\mathrm{lat}}}{p_x}. \tag{S39}$$

Finally the whole volume is rotated by $\theta$ about the $x$ axis using,

$$\begin{bmatrix} 1 & 0 & 0 \\ 0 & \cos\theta & -\sin\theta \\ 0 & \sin\theta & \cos\theta \end{bmatrix} \tag{S40}$$

where, the sheet angle, $\theta$, is given by,

$$\theta = \arctan\left(\tan\theta' \frac{n_{\text{sample}}}{M_{\text{lat}}}\right) \tag{S41}$$

where, $n_{\text{sample}}$ is the refractive index of the sample. This renders the volume in lab coordinates and the resultant $xy$ stack is equivalent to that obtained from a standard confocal microscope. Combining these transformations into a single operation we get,

$$\begin{aligned}\begin{bmatrix}x\\y\\z\end{bmatrix} &= \begin{bmatrix}1&0&0\\0&\cos\theta&-\sin\theta\\0&\sin\theta&\cos\theta\end{bmatrix}\begin{bmatrix}1&0&0\\0&1&0\\0&0&\epsilon\end{bmatrix}\begin{bmatrix}1&0&0\\0&1&\gamma\\0&0&1\end{bmatrix}\begin{bmatrix}1&0&0\\0&S_y&0\\0&0&1\end{bmatrix}\begin{bmatrix}x'\\y'\\z'\end{bmatrix}\\ &= \begin{bmatrix}1&0&0\\0&S_y\cos\theta&\gamma\cos\theta-\epsilon\sin\theta\\0&S_y\sin\theta&\gamma\sin\theta+\epsilon\cos\theta\end{bmatrix}\begin{bmatrix}x'\\y'\\z'\end{bmatrix}.\end{aligned} \tag{S42}$$

This renders a transformation from the raw coordinate frame $(x', y', z')$ to the lab coordinate frame $(x, y, z)$ of,

$$\begin{aligned}x &= x',\\ y &= y'S_y\cos\theta + z'\left(\gamma\cos\theta - \epsilon\sin\theta\right),\\ z &= y'S_y\sin\theta + z'\left(\gamma\sin\theta - \epsilon\cos\theta\right).\end{aligned} \tag{S43}$$

Similarly, by calculating the inverse of equation S42 we can obtain the transformation from the lab coordinate frame back to the raw coordinate frame using,

$$\begin{bmatrix}x'\\y'\\z'\end{bmatrix} = \begin{bmatrix}1&0&0\\0&\frac{\gamma\sin\theta+\epsilon\cos\theta}{S_y\epsilon}&\frac{\epsilon\sin\theta-\gamma\cos\theta}{S_y\epsilon}\\0&-\frac{\sin\theta}{\epsilon}&\frac{\cos\theta}{\epsilon}\end{bmatrix}\begin{bmatrix}x\\y\\z\end{bmatrix} \tag{S44}$$

and

$$\begin{aligned}x' &= x,\\ y' &= \frac{y\left(\gamma\sin\theta+\epsilon\cos\theta\right)+z\left(\epsilon\sin\theta-\gamma\cos\theta\right)}{S_y\epsilon},\\ z' &= \frac{z\cos\theta - y\sin\theta}{\epsilon}.\end{aligned} \tag{S45}$$

In this work we perform deskewing using a custom Python script that loops through a volume in the lab coordinate frame $(x, y, z)$ and uses the transformation provided by equation S45 to find the pixel in the raw data that corresponds to that location. When this does not line up with an exact pixel, linear interpolation is used to calculate the appropriate pixel value. This method has the benefit of being memory efficient and being able to deskew large datasets on a standard workstation. During deskewing, algorithms for hot pixel correction and illumination flatness correction are also applied.

### F Resolution enhancement via sub-pixel image shifting

Due to the size of the camera pixels in CtDvOPM the raw data is undersampled, leading to a Nyquist sampling resolution limit of 3.77 µm. In order to fully exploit the diffraction limited imaging performance of the 0.125 NA bi-telecentric lens used for imaging, we employ a beam shifter to shift the image on the camera by half a pixel along both camera axes. This allows the reconstruction of images that are equivalent to having acquired the data with a camera with pixels of half the size. In this case, a Nyquist sampling limit of 1.88 µm (0.94 µm pixels in the sample) is achieved, enabling sampling slightly beyond the diffraction limited resolution of 2.0 µm. In this section we present a comparison of the algorithms used to reconstruct upsampled data from a set sub-pixel shifted images (Supplementary Figure 5) as well as two examples of sub-pixel shifting applied to imaging a SmartBatch+ cleared mouse brain (Supplementary Figure 6) and a CUBIC cleared mouse cerebellum (Supplementary Figure 7). In these examples we demonstrate how the image shifting upsampling allows structures to be identified that cannot be resolved in the raw or bicubic interpolated data.

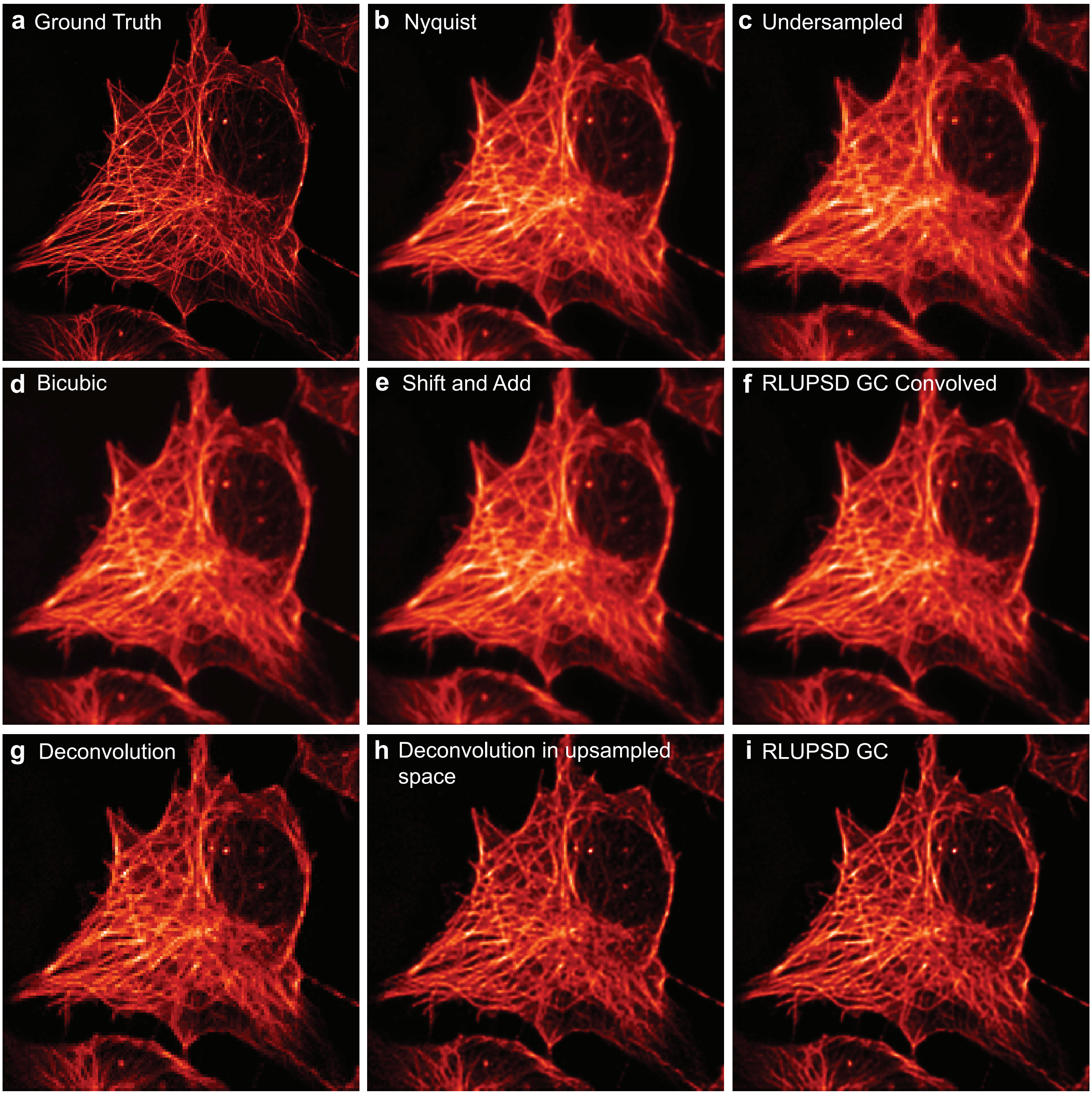


**Supplementary Figure 5: Comparision of algorithms for reconstructing upsampled data using simulated ground truth data.**
**a** Ground truth data used for generating the simulated data. **b** Ground truth data convolved with a simulated point spread function (PSF) and sampled at the Nyquist limit. **c** Image of b binned 2 × 2, to match raw experimental data. **d** Bicubic interpolation of a single undersampled raw frame c. **e** Upsampled reconstruction using four sub-pixel shifted raw frames using the shift and add algorithm. **f** Upsampled reconstruction using four sub-pixel shifted raw frames using a Richardson–Lucy combined image shifting and deconvolution algorithm, with a gradient consensus stopping criterion, where the final result is convolved with the system PSF (i.e. the final result is not deconvolved, only upsampled). **g** Result of a Richardson–Lucy deconvolution of an undersampled raw frame using an undersampled system PSF. Note that microtubules in the bottom left of the cell are incorrectly shown to have a criss-cross structure. **h** Result of a Richardson–Lucy deconvolution of an undersampled raw frame into an upsampled grid using a Nyquist sampled system PSF. Note that the erroneous criss-cross structure shown in g is still present although the pixel grid is finer. **i** Repeat of f omitting the final convolution step, with no visible criss-cross error.

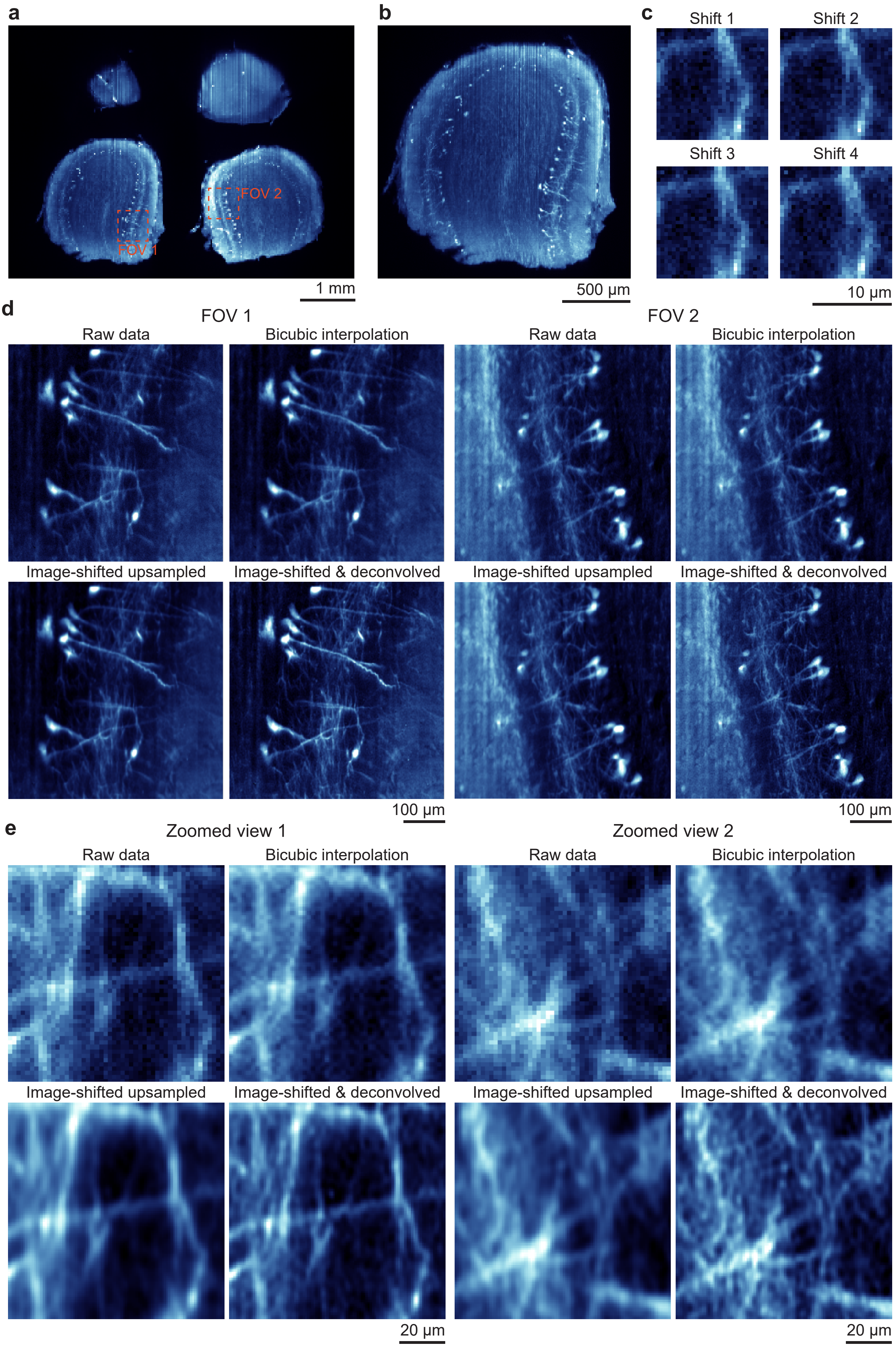

a
FOV 1
FOV 2
1 mm
b
500 µm
c
Shift 1
Shift 2
Shift 3
Shift 4
10 µm
d
FOV 1
FOV 2
Raw data
Bicubic interpolation
Raw data
Bicubic interpolation
Image-shifted upsampled
Image-shifted & deconvolved
Image-shifted upsampled
Image-shifted & deconvolved
100 µm
100 µm
e
Zoomed view 1
Zoomed view 2
Raw data
Bicubic interpolation
Raw data
Bicubic interpolation
Image-shifted upsampled
Image-shifted & deconvolved
Image-shifted upsampled
Image-shifted & deconvolved
20 µm
20 µm

**Supplementary Figure 6: Exemplar data of a image-shifted upsampled SmartBatch+ cleared mouse brain.**
**a** Raw frame of SmartBatch+ cleared mouse brain indicating the location of the two FOVs presented later in the figure. **b** Cropped region of a. **c** Exemplar images of 2 × 2 sub-pixel shifted images of the same FOV. **d** Comparison between raw, bicubic interpolated, image-shifted upsampled and deconvolved image-shifted upsampled data of two FOV shown in a. **e** Zoomed view of the data shown in d clearly demonstrating the increase in image quality of the image-shifted upsampled data.

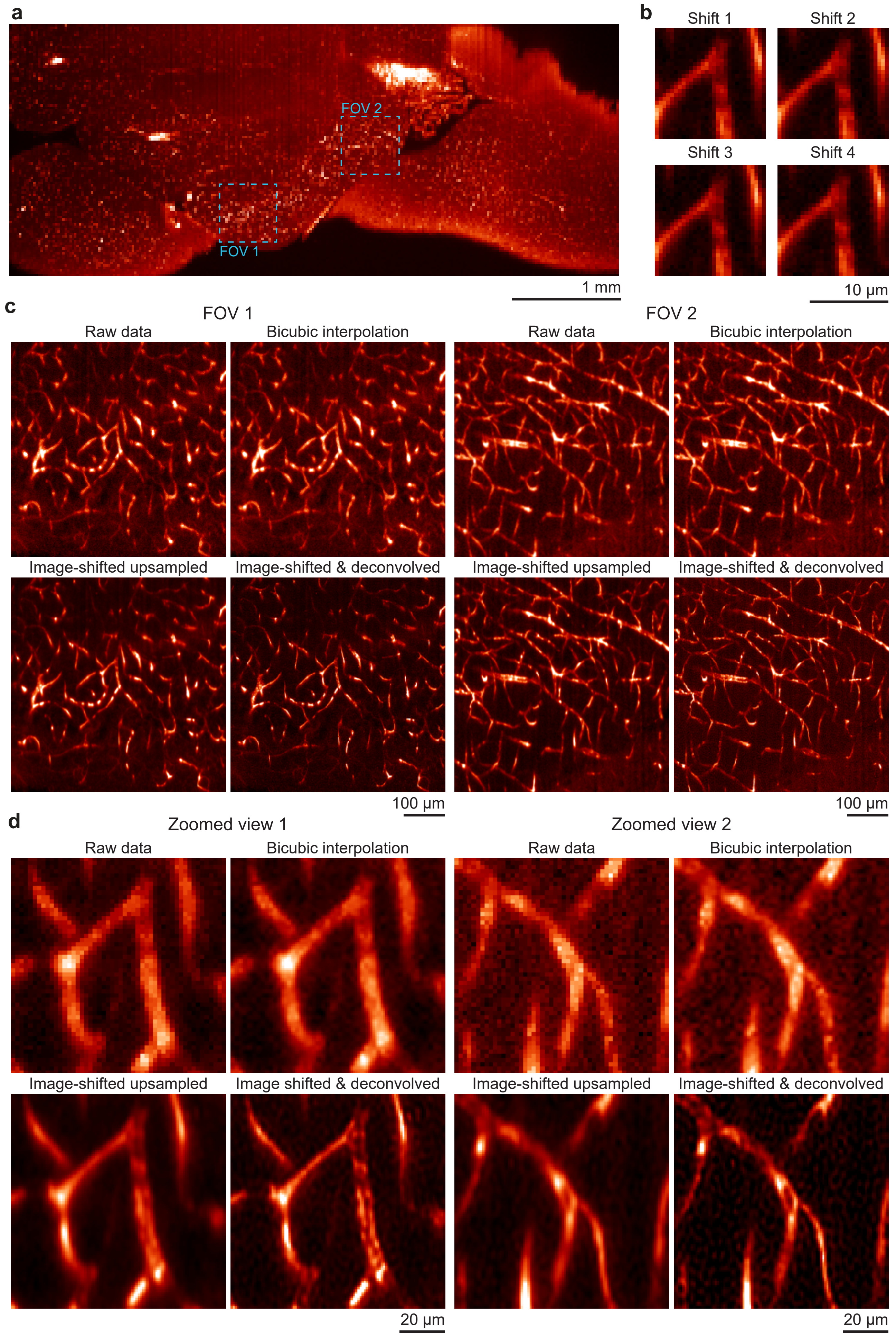
a
FOV 2
FOV 1
1 mm
b
Shift 1
Shift 2
Shift 3
Shift 4
10 μm
c
FOV 1
Raw data
Bicubic interpolation
Image-shifted upsampled
Image-shifted & deconvolved
100 μm
FOV 2
Raw data
Bicubic interpolation
Image-shifted upsampled
Image-shifted & deconvolved
100 μm
d
Zoomed view 1
Raw data
Bicubic interpolation
Image-shifted upsampled
Image shifted & deconvolved
20 μm
Zoomed view 2
Raw data
Bicubic interpolation
Image-shifted upsampled
Image-shifted & deconvolved
20 μm

**Supplementary Figure 7: Exemplar data of a image-shifted upsampled CUBIC cleared mouse cerebellum.**
**a** Raw frame of CUBIC cleared mouse cerebellum indicating the location of the two FOVs presented later in the figure. **b** Exemplar images of $2 \times 2$ sub-pixel shifted images of the same FOV. **c** Comparison between raw, bicubic interpolated, image-shifted upsampled and deconvolved image-shifted upsampled data of two FOV shown in a. **d** Zoomed view of the data shown in d clearly demonstrating the ability visualise structures in the image-shifted upsampled data that is unresolvable in the raw or bicubic interpolated data (also see Figure 3e).
